# Biogenesis and Function of *c-*type Cytochromes in the Methanogenic Archaeon, *Methanosarcina acetivorans*

**DOI:** 10.1101/2022.01.26.477811

**Authors:** Dinesh Gupta, Katie E. Shalvarjian, Dipti D. Nayak

## Abstract

*C*-type cytochromes (cyt *c*) are proteins that covalently bind heme and are integral to electron transport chains. A growing body of evidence suggests that cyt *c* play a vital role in both intra- and extra-cellular electron transfer processes in Archaea, especially in members that metabolize methane and other short chain alkanes. Elaborate mechanisms for the biogenesis of cyt *c* are known in Bacteria and Eukarya but this process remains largely uncharacterized in Archaea. Here, we have used the model methanogenic archaeon *Methanosarcina acetivorans* to characterize a distinct form of the system I cyt *c* maturation machinery (referred to as the Ccm machinery henceforth) that is broadly distributed in members of the Archaea. Phenotypic analyses of *M. acetivorans* mutants deficient in essential components of the Ccm machinery reveal that cyt *c* are broadly important for growth and methanogenesis, but the magnitude of their impact can vary substantially depending on the growth substrate. Heterologous expression of a synthetic operon with the Ccm machinery (CcmABCEF) from *M. acetivorans* is both necessary and sufficient for cyt *c* biogenesis in a non-native host (*M. barkeri* Fusaro) that is incapable of cyt *c* biogenesis. Even though components of the Ccm machinery are universally conserved across the Archaea, our evolutionary analyses indicate that different clades of Archaea acquired this pathway through multiple independent horizontal gene transfer events from different groups of Bacteria. Overall, we have demonstrated the convergent evolution of a novel Archaea-specific Ccm machinery for cyt *c* biogenesis and its role in methane metabolism.

**Significance Statement:** Microorganisms belonging to the domain Archaea play an especially important role in regulating atmospheric methane levels. Specifically, methanogens are the primary source of biogenic methane and **<u>an</u>**aerobic **<u>me</u>**thanotrophic archaea (ANME) consume a substantial proportion of methane released in marine sediments. Genomic studies have implicated a class of electron-transfer proteins called *c*-type cytochromes as being crucial in mediating archaeal methane metabolism in the environment. However, neither the biogenesis nor the role of *c*-type cytochromes in methane metabolism has ever been investigated. Here, we have used a model methanogen, *Methanosarcina acetivorans,* to characterize a distinct pathway for maturation of *c*-type cytochromes that seems to be uniformly conserved across the Archaea and have also identified substrate-specific functional roles for *c*-type cytochromes during methanogenesis.

## Introduction

*C*-type cytochromes [Cyt(s) *c*] are found across the tree of Life, and are critical to electron transfer processes ranging from aerobic respiration in the mitochondrion of Eukaryotes to intra-/extra-cellular electron transport in Bacteria and Archaea [1–4]. The production of a holo-cyt *c*, that is capable of electron transfer, requires complex post-translational processing to covalently attach iron-protoporphyrin IX (heme *b*) to an unfolded apo-cyt *c* protein [5, 6]. Heme attachment occurs on the extracellular face of the cytoplasmic membrane, and, thus also requires the translocation of the apoprotein and heme across the membrane. The apoprotein is translocated via the Sec pathway [7–9] and a set of proteins collectively known as the cyt *c* maturation system are involved in energy-dependent heme transport, apoprotein handling, and covalent cofactor attachment [5, 6, 10]. Three distinct systems for cyt *c* maturation have been identified thus far, and have been studied extensively in bacterial and eukaryotic model systems [5, 6]. However, despite growing evidence that cyt *c* play a vital role in electron transport processes across members of the Archaea - particularly in microorganisms that metabolize methane and other alkanes- the biogenesis of cyt *c* has never been functionally characterized in an archaeon.

Based on genomic surveys [11–13] of sequenced isolates and metagenomics assembled genomes (MAGs), the vast majority of cyt *c* containing Archaea encode homologs of the System I cyt *c* maturation pathway (Ccm pathway) that is widespread in members of the Bacteria and is also present in the mitochondria of some Plants and Protozoa [5, 14]. The Ccm pathway is best characterized in Gram-negative bacteria, such as *E. coli,* where nine to ten proteins, encoded by *ccmABCDEFGH*(*I*) and *ccdA* (or *dsbD*), are involved in cyt *c* biogenesis (Supplementary Figure 1) [5, 6]. Typically, the *ccm* genes are encoded in an operon on the chromosome and the *ccmH* open reading frame in *E. coli* is often found as two different genes, annotated as *ccmH* (or *ccl2*) and *ccmI* (or *cycH*), in bacteria like *Rhodobacter capsulatus* [5]. Genetic and biochemical studies of the Ccm pathway have revealed that the membrane-associated CcmABCD complex translocates heme across the cytoplasmic membrane to form holo-CcmE (Supplementary Figure 1). Heme from holo-CcmE is ultimately transferred to the apo-cyt *c* by the CcmF/H complex (cytochrome synthetase) to form holo-cyt *c*. CcmG and CcdA (or DsbD) reduce the disulfide bond between cysteine residues of heme-binding motifs (CXXCH) in the apo-cyt *c* prior to heme attachment (Supplementary Figure 1)[15–17]. Curiously, most, if not all, archaeal genomes sequenced thus far lack several *ccm* genes (namely *ccmD, ccmH, ccmI)* that have been shown to be essential for cyt *c* biogenesis in Bacteria like *E. coli, Rhodobacter capsulatus, Paracoccus denitrificans, Bradyrhizobium japonicum, Shewenella oneidensis,* and *Desulfovibrio desulfuricans* [10, 18–20]. In bacteria, CcmD facilitates the release of the heme-bound holo-CcmE from the CcmABCD complex [21], while CcmH and CcmI are a part of the protein complex involved in covalent attachment of heme to the apo-cyt *c* (Supplementary Figure 1)[5, 6]. As such, based on evidence from studies with bacteria, the streamlined Ccm machinery found in archaeal genomes, comprised only of CcmABCEFG and CcdA, would be insufficient for the biogenesis of cyt *c*. However, the mature cyt *c* proteins have been identified in many archaeal strains, such as *Haloferax volcanii, Ignicoccus hospitalis, Pyrobaculum islandicum, Ferroglobus placidus,* and *Methanosarcina spp* [12, 22–26]. Whether Archaea have replaced CcmD, CcmH, and CcmI with non-orthologous proteins or have reconfigured the Ccm machinery such that *ccmDHI* are no longer essential remains unclear.

In this study, we used the genetically tractable methanogenic archaeon, *Methanosarcina acetivorans,* as a model system to functionally characterize the pathway for biogenesis of cyt *c* in archaea. Most archaea that encode cyt *c* proteins are either recalcitrant to laboratory cultivation techniques, genetically intractable, or only encode only one cyt *c* that might be essential for growth [11–13, 27]. In contrast, *M. acetivorans* encodes multiple cyt *c* proteins [28], can be easily cultivated in a laboratory [29], and has state-of-the-art genetic tools [30, 31], including inducible gene expression, tests for gene essentiality, and CRISPR-Cas9 based genome editing for complex genetic manipulation [32]. *M. acetivorans* is a methanogen that can only grow by coupling energy conservation to the production of methane gas using acetate or methylated compounds, like methanol, = as a growth substrate. At least 5 different cyt *c* proteins are encoded in the *M. acetivorans* genome and these proteins contain between 1 and 7 heme-binding motifs [28]. Recent studies have even proposed distinct roles for some of these cyt *c* proteins in intracellular electron transport during methanogenesis [26, 33–35], **<u>e</u>**xtracellular **<u>e</u>**lectron **t**ransfer (EET) [28] as well as **<u>d</u>**irect **i**nterdomain **<u>e</u>**lectron **<u>t</u>**ransfer (DIET) between *M. acetivorans* and bacterial strains like *Geobacter metallireducens* [36]. Thus, *M. acetivorans* is not only an ideal candidate to dissect the biogenesis of cyt *c* in Archaea but also to functionally characterize the role of cyt *c* in various electron transfer processes that have ramifications on global carbon cycling as well as on the formation of microbial communities in anoxic environments.

Here, we have used a combination of genetic, molecular, and biochemical analyses, to show that *M. acetivorans* and other archaea use a streamlined version of the Ccm machinery that only requires *ccmABCEF* for cyt *c* biogenesis. To this end, we have shown that the *ccmABCEF* from *M. acetivorans* is sufficient to produce holo-cyt *c* in a heterologous host, *Methanosarcina barkeri* Fusaro, that otherwise is incapable of cyt *c* biogenesis. Our physiological analyses also reveal substrate-specific phenotypes for the cyt *c* biogenesis pathway and cyt(s) *c* during growth and methanogenesis in *M. acetivorans*. A closer inspection of the distribution and synteny of the cyt *c* biogenesis genes in methane-metabolizing archaea related to *Methanosarcina* (i.e. belonging to the Order *Methanosarcinales*) suggests that the cyt *c* biogenesis genes were likely acquired in the last common ancestor of the *Methanosarcinales* and have been lost in many extant clades that also do not encode any cyt *c* genes. Although the streamlined Ccm machinery is conserved across Archaea, our evolutionary analyses suggest that the acquisition of cyt *c* biogenesis in Archaea has occurred through multiple horizontal gene transfer (HGT) events with different members of the Bacteria. Overall, we have used the model methanogenic archaeon, *M. acetivorans,* as a model system to characterize a streamlined form of the Ccm machinery used for the biogenesis of cyt *c* in members of Archaea.

## Results

### The Ccm machinery is essential for *c*-type cytochrome biogenesis in *M. acetivorans*

We used a sequence-based approach to identify homologs of eight different genes of the Ccm machinery for cyt *c* maturation at four different chromosomal loci in *M. acetivorans.* The *ccmABC* genes (MA1428-MA1430) are located at one chromosomal locus and the *ccmE* gene (MA4149) is in a putative operon with a geranyl farensyl diphosphate synthase at another locus (Supplementary Figure 1). Two *ccmF* genes (MA3305 and MA3304 that have been renamed *ccmF1* and *ccmF2* respectively), which likely represent the C- and N- terminal segments of the bacterial *ccmF* locus respectively, are encoded in a putative operon. Finally, *ccmG* and *ccdA* are found adjacent to each other on the chromosome (MA4254 and MA4255) (Supplementary Figure 1). We used our recently developed Cas9-based genome editing technology to generate markerless in-frame deletion mutants of the *ccmABC* locus, *ccmE*, *ccmF1, ccmF2*, *ccmG, ccdA* as well as the Δ*ccmF1ΔccmF2* and Δ*ccmG*Δ*ccdA* double mutants. We picked the Δ*ccmABC* mutant for whole-genome sequencing to screen for off-target CRISPR-Cas9 activity. Apart from the deleted locus, we did not observe any other mutations in the Δ*ccmABC* mutant relative to the parent strain (WWM60) (Supplementary Table 1). Consistent with our previous observations, these data suggest that off-target activity is negligible during Cas9-mediated genome editing in *M. acetivorans* [32, 37, 38].

We selected a representative cyt *c* encoded by the *mmcA* gene (locus tag: MA0658) in the *M. acetivorans* genome to assay for cyt *c* maturation in the *ccm* mutants. MmcA is a heptaheme cyt *c* that contains a N-terminal signal peptide and is likely located in the pseudo-periplasm (Supplementary Figure 2). MmcA is a highly expressed cyt *c* that is conserved across methane-metabolizing archaea [28, 39, 40] and holo-MmcA is easy to detect as it contains seven heme-binding motifs. Taken together, these functional and technical features make MmcA an ideal candidate to assay cyt *c* biogenesis. To develop a rapid assay for quantification of holo-cyt *c* in the *ccm* mutants, we built a plasmid-based overexpression system by adding a C-terminal TAP (**<u>t</u>**andem **<u>a</u>**ffinity **<u>p</u>**urification) tag comprised of a 3X FLAG sequence and a twin-Strep sequence to the *mmcA* coding sequence placed under the control of a tetracycline-inducible promoter (Figure 1a). As a control, we transformed the Δ*mmcA* mutant with an untagged or TAP-tagged cyt *c* overexpression vector to test if the presence of the tag in the MmcA CDS interferes with protein production or heme-attachment. To identify protein production, we performed an immunoblot with a commercial anti-Flag antibody. Using this technique, we were able to successfully detect the TAP-tagged MmcA in the cell lysate (Figure 1b). To assay for heme attachment (formation of holo-cyt *c*), we used a peroxidase-based heme stain, which detects covalently bound (*c*-type) heme in cyt *c* [41]. Using this technique, we were able to verify that the TAP-tagged MmcA is capable of undergoing covalent modification to generate the corresponding holo cyt *c* (Figure 1b). Taken together, these controls demonstrate that our overexpression vectors can be used to reliably and rapidly assay the production and covalent modification of MmcA in *M. acetivorans*.

**Figure 1:**
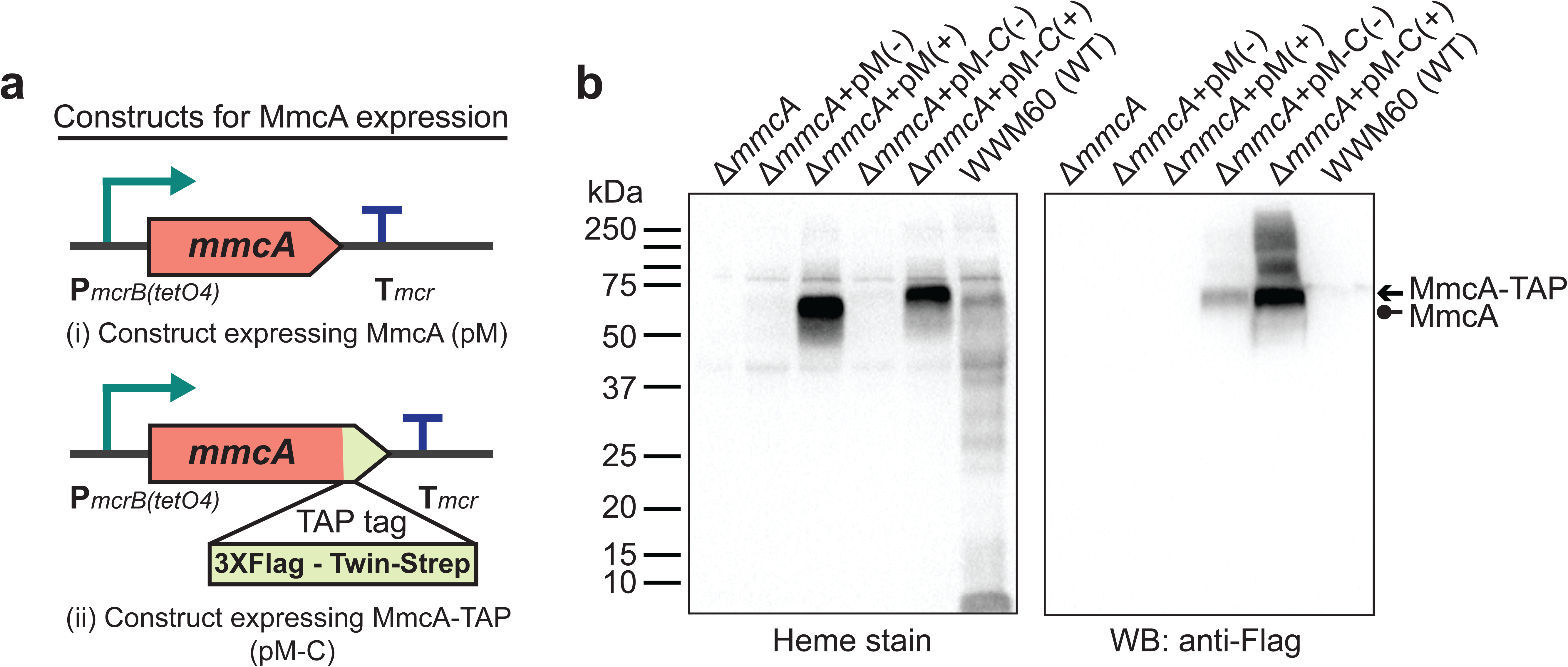
**a)** The design of diagnostic constructs to use the native heptaheme *c*-type cytochrome, MmcA, to map the cytochrome *c* biogenesis pathway in *Methanosarcina acetivorans. i)* The control construct (pM) contains a *mmcA* coding sequence whereas *ii)* the test construct (pM-C) contains a *mmcA* coding sequence with a C-terminal translational fusion of a tandem affinity purification (TAP) tag comprised of a 3X FLAG sequence and a Twin-Strep sequence. In each construct, the gene expression driven by a tetracycline-inducible medium-strength promoter, P*mcrB*(*tetO4*), and a transcriptional terminator of the *mcr* operon from *M. acetivorans* is provided after the last coding sequence. **b)** Assays to measure holo-MmcA formation by covalent heme attachment and detect the tagged MmcA protein in whole cell lysates of *M. acetivorans* mutants. (*Left*) A heme peroxidase-based assay is used to detect the presence of proteins with covalently bound heme in whole cells lysates. Using this assay, both untagged (Lane 3) and tagged (Lane 5) heme-bound holo-MmcA can be detected upon induction of the genes from each plasmid construct described above. The tagged MmcA protein runs at a higher molecular weight due to the presence of a *ca.* 7.3 kDa TAP tag at the C-terminus compared to the native MmcA protein. (*Right*) Immunoblotting with commercial anti-FLAG antibody can be used detect MmcA production in the test construct containing a C-terminal TAP tag fused to the *mmcA* gene (Lanes 4 and 5). (-) indicates that no tetracycline was added to the growth medium and (+) indicates that 100 µg/ml tetracycline was added to the growth medium. An equal amount of whole cell lysate protein (∼80 µg) was loaded in each lane.

Next, we introduced the plasmid overexpression system with a TAP-tagged MmcA in each of the *ccm* deletion mutants and assayed for the production of tagged MmcA protein and holo-MmcA in each of the resulting strains (Figures 2a-2c). We were unable to detect the tagged protein or holo-MmcA in the Δ*ccmABC, ΔccmE, ΔccmF2, ΔccmF1ccmF2* mutants, which suggests that these genes are essential for cyt *c* biogenesis in *M. acetivorans* (Figure 2c). Consistent with previous studies, these results also indicate that the apo-MmcA is immediately targeted for degradation in the absence of a functional Ccm machinery [42–44]. Curiously, we were able to detect faint band corresponding to holo-MmcA in the Δ*ccmF1* mutant (Figure 2c). These data suggest that the gene product of *ccmF1* is important but not vital for cyt *c* biogenesis possibly because the catalytically important residues from CcmF in *E. coli* are all present in CcmF2 from *M. acetivorans* (Supplementary Figure 3). Finally, we observed that Δ*ccmG* and Δ*ccdA* single mutants as well as the Δ*ccmG*Δ*ccdA* double mutant produced roughly the same amount of holo-MmcA as the parent strain (Figure 2c), which indicates that these genes are not essential for disulfide bond reduction in the apo cyt *c* under highly reducing laboratory growth conditions. Overall, using MmcA as a diagnostic cyt *c*, we have shown that the proteins encoded by *ccmABCEF1F2* in *M. acetivorans* constitute a functional, streamlined version of the Ccm machinery for cyt *c* maturation.

**Figure 2:**
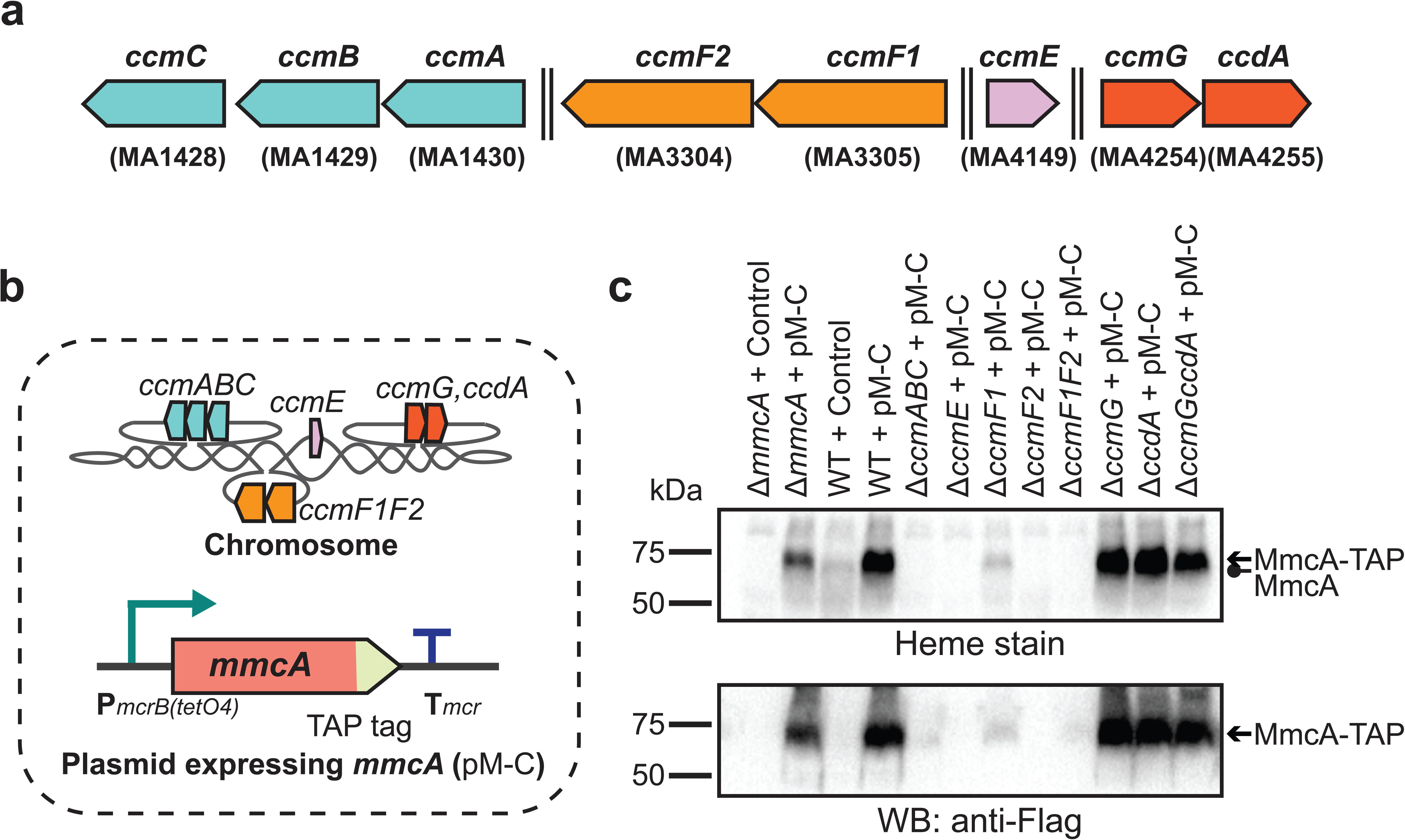
**a)** Chromosomal organization of the system I cytochrome *c* maturation machinery (Ccm) genes in *M. acetivorans.* Double dashed lines indicate that the genes are located more than 110 kbp away on the chromosome. Genes located at the same chromosomal locus are shown in the same color. **b)** A schematic showing the genotype of strains used to assay the role of individual *ccm* genes in the maturation of *c*-type cytochromes. In addition to the corresponding *ccm* deletion on the chromosome, these strains also contain a self-replicating plasmid (pM-C) where the *mmcA* coding sequence with a C-terminal tandem affinity purification (TAP) tag comprised of a 3X FLAG sequence and a Twin-Strep sequence is placed under the control of a tetracycline-inducible medium-strength promoter, P*mcrB*(*tetO4*). In these strains, the expression of the C-terminal TAP tagged MmcA can be induced by the addition of tetracycline to the growth medium. **c)** A heme peroxidase-based assay to measure the formation of heme-bound holo-MmcA (top) and Western blots (WB) with anti-FLAG antibody to detect the production of C-terminal TAP tagged MmcA (bottom). All assays were conducted with whole cell lysates of cultures grown in medium containing 100 µg/ml tetracycline to induce the expression of C-terminal TAP tagged MmcA. Neither holo-MmcA nor tagged protein could be detected in the Δ*ccmCBA*, Δ*ccmE*, Δ*ccmF2* single mutants and the Δ*ccmF1*Δ*ccmF2* double mutant suggesting that the apo-MmcA undergoes rapid proteolysis in the absence of a functional Ccm machinery. An equal amount of whole cell lysate protein (∼60 µg) was loaded in each lane. The vector control used for this experiment is pJK029A as described previously elsewhere (Guss *et al*., 2008).

### Archaeal CcmABC is involved in the formation of a holo-CcmE heme chaperone

In the first steps of cyt *c* biogenesis, heme *b* is transported across the cytoplasmic membrane to generate holo-CcmE: a heme chaperone that covalently binds heme and transports it to the cytochrome *c* synthetase complex [45, 46]. In Gram-negative bacteria, like *E. coli*, heme transport and holo-CcmE formation is mediated by the CcmABCD complex (Figure 3a). In the CcmABCD complex, CcmD plays an essential role in facilitating the release of holo-CcmE from the CcmABCE adduct [5, 21]. Since *M. acetivorans* and all other sequenced archaeal strains lack CcmD, we investigated the role of the CcmABC complex in the formation of holo-CcmE (Figure 3b). To this end, we developed a plasmid-based overexpression system for CcmE with a C-terminal 1XStrep-1XFlag tag placed under the control of a tetracycline-inducible promoter (Figure 3c). We introduced the C-tagged CcmE overexpression plasmid in the parent strain (WWM60) as well as the Δc*cmABC* and Δ*ccmE* mutants and were able to detect tagged-CcmE in protein enriched from the membrane fraction (using an anti-Strep affinity column) by immunoblotting with an anti-FLAG antibody in all plasmid complemented strains (Figure 3d). We were only able to observe the heme-bound holo-CcmE by heme staining the enriched membrane fraction of the plasmid complemented parent strain (WWM60) and Δ*ccmE* mutant (Figure 3d). Furthermore, we were unable to detect holo-cyt *c,* including the MmcA protein, by heme staining the total cell lysate of the Δ*ccmABC* strain complemented with the CcmE overexpression plasmid, which indicates that overexpression of CcmE does not rescue the *ccmABC* lesion in the cytochrome maturation machinery (Supplementary Figure 4). These data support the hypothesis that the CcmABC complex has evolved to transport heme and produce holo-CcmE independent of CcmD in *M. acetivorans* and other archaea.

**Figure 3:**
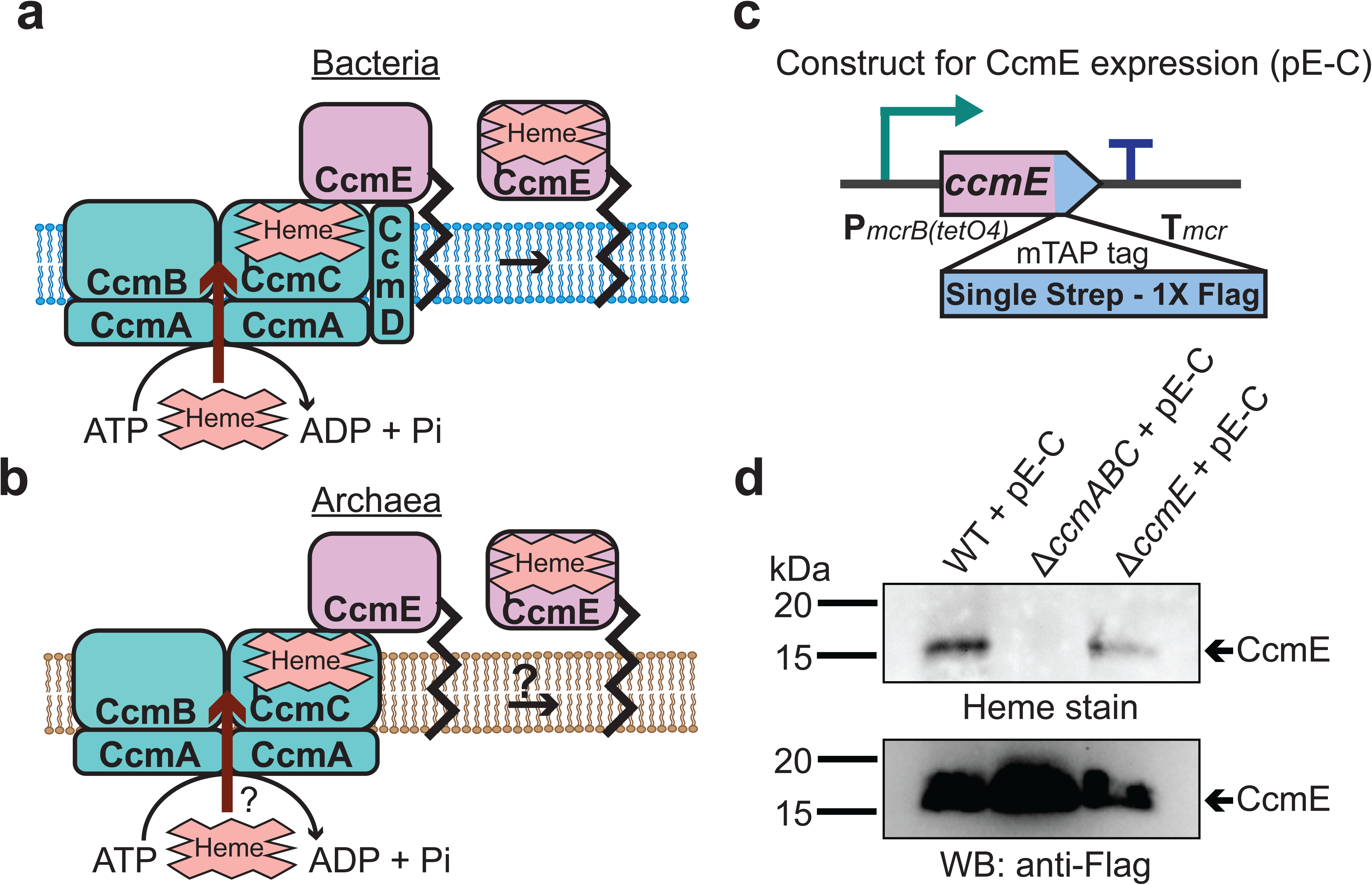
**a)** The formation of heme-bound holo-CcmE in bacteria containing the System I Ccm machinery is facilitated by the CcmABCD complex. The ATP-dependent CcmABC complex translocates heme *b* from the cytosol to CcmE and CcmD facilitates the dissociation of holo-CcmE from the CcmABCDE adduct. **b)** Since archaea lack the *ccmD* gene the formation of holo-CcmE is unclear. One hypothesis, shown here, is that the CcmABC complex in archaea mediates the dissociation of holo-ccmE independent of CcmD. **c)** The design of a test construct (pE-C) containing the *ccmE* coding sequence with C-terminal translational fusion of a modified tandem affinity purification tag (mTAP) comprising of a Single-Strep sequence and a 1XFLAG sequence. In this construct, the coding sequence is placed in between a tetracycline-inducible medium-strength promoter, P*mcrB*(*tetO4*) (ref), and the transcriptional terminator of the *mcr* operon from *M. acetivorans.* Here, the expression of the C-terminal mTAP tagged CcmE can be induced by the addition of tetracycline to the growth medium. **d)** A heme peroxidase-based assay to measure the formation of holo-CcmE (top) and a Western blot (WB) with anti-FLAG antibody to detect the production of C-terminal mTAP tagged CcmE (bottom) was conducted. Even though CcmE is produced in the Δ*ccmABC* mutant, holo-CcmE cannot be formed, consistent with the model shown in **b)**. All assays were performed with an enriched membrane fraction (i.e. protein eluted after passing the membrane fraction through a Strep-affinity column) of cultures grown in medium containing 100 µg/ml tetracycline to induce the expression of C-terminal mTAP tagged CcmE. An equal amount of protein (∼1 µg) was loaded in each lane.

### A CXXXY motif in archaeal CcmE is required for heme-attachment and protein stability

In bacteria with a Ccm machinery, like *E. coli,* a conserved Histidine residue (H130) present in the HXXXY motif of CcmE covalently binds heme [47]. Notably, all sequenced archaea (and a few bacteria like *Desulfovibrio desulfuricans)* have replaced this Histidine with a Cysteine (Figure 4a) [11, 19]. To test if the Cysteine residue (C120) in the conserved motif (CXXXY) of CcmE in *M. acetivorans* has a functional role similar to the Histidine residue in *E. coli* or the Cysteine residue in *D. desulfuricans*, we built a plasmid-based overexpression system for mutant alleles of *ccmE* with either a C120A or a C120H substitution and a C-terminal 1XStrep-1XFlag tag under the control of an inducible promoter (Figures 4a and 4b). We introduced these plasmids in the Δ*ccmE* mutant and successfully detected the wild-type and C120A CcmE protein in the Strep-enriched membrane fraction by immunoblotting with anti-FLAG antibodies (Figure 4c). Despite repeated attempts, we were unable to detect the C120H CcmE mutant in the membrane or soluble fraction (Figure 4c and Supplementary Figure 5). This outcome is consistent with the hypothesis that the C120H substitution considerably destabilizes CcmE, leading to proteolysis of the gene product. Next, we were able to observe the wild-type CcmE by heme staining the Strep-enriched membrane fraction of the corresponding strain however, no heme stained band corresponding to CcmE was detected for the C120A mutant (Figure 4c). Furthermore, we were unable to detect holo-MmcA or any other holo-cyt *c* by heme staining the total cell lysate of the Δ*ccmE* strain complemented with either the C120A or C120H CcmE (Supplementary Figure 6). These results provide strong evidence in support of the hypothesis that the C120 residue of the CXXXY motif is important for heme attachment as well as protein stability in the archaeal CcmE.

**Figure 4:**
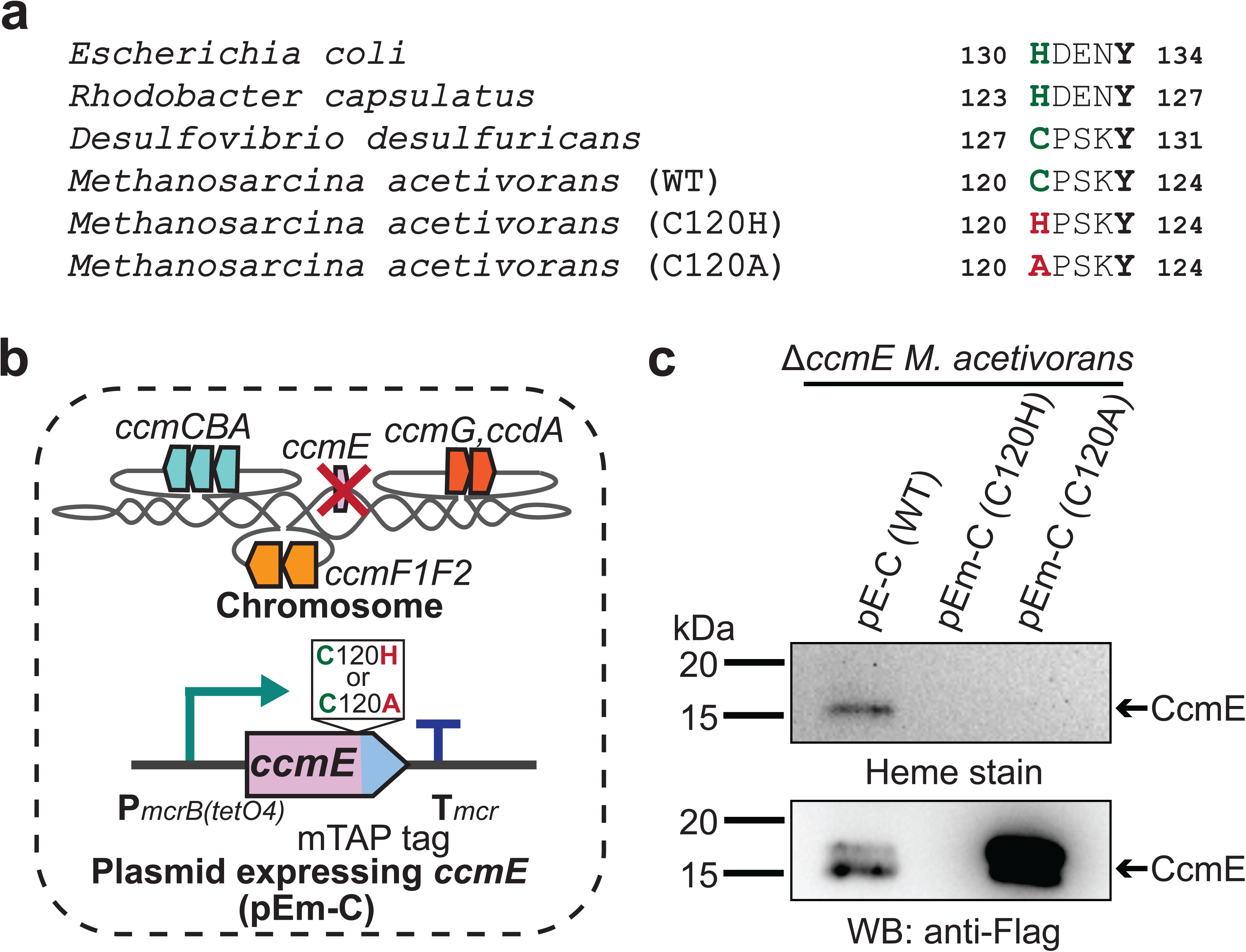
**a)** Alignment of the heme-binding domain in CcmE sequences derived from representative bacteria and archaea. The heme binding residue in bacteria can vary: some species like *E. coli* covalently bind heme using a histidine residue whereas others like *Desulfovibrio desulfuricans* covalently bind heme using a cysteine residue. Almost all archaeal CcmE sequences, including the sequence derived from *M. acetivorans,* contain a conserved cysteine residue that is likely involved in heme binding. **b)** To test the role of the cysteine residue in the CcmE sequence derived from *M. acetivorans,* we expressed the C120A and C120H point mutants of *ccmE* with a modified tandem affinity purification tag (mTAP) comprising of a Single-Strep sequence and a 1XFLAG sequence at the C-terminus as described in Figure 3c. **c)** A heme peroxidase-based assay was used to measure the formation of holo-CcmE (top) and Western blot (WB) with anti-FLAG antibody was used to detect the production of C-terminal mTAP tagged CcmE (bottom). Neither the C120A or the C120H mutants of CcmE contain covalently bound heme (top); furthermore, no mTAP-tagged CcmE was detected by WB for the C120H mutant (bottom), likely indicating that this point mutation destabilizes the protein substantially. All assays were performed with an enriched membrane fraction (i.e. protein eluted after passing the membrane fraction through a Strep-affinity column) of cultures grown in medium containing 100 µg/ml tetracycline to induce the expression of C-terminal mTAP tagged CcmE. An equal amount of protein (∼1 µg) was loaded in each lane.

### A streamlined Ccm machinery comprised of CcmABCEF is necessary and sufficient for *c*-type cytochrome biogenesis in *Methanosarcina spp*

While our genetic studies with *M. acetivorans* clearly show that the proteins encoded by *ccmABC, ccmE, ccmF1* and *ccmF2* are essential for cyt *c* biogenesis, they do not inform us of any non-orthologous proteins that might also be involved in this process. To test if the cyt *c* maturation pathway encoded by the *ccm* genes in *M. acetivorans* is both necessary and sufficient for cyt *c* biogenesis, we expressed *mmcA* and a synthetic operon comprised of *ccmABCEF1F2* in a heterologous host. We chose *Methanosarcina barkeri* Fusaro as a heterologous host for this study as it has the same codon usage pattern as *M. acetivorans*, contains an intact pathway for heme *b* synthesis, and is known to produce *b*-type cytochromes, but does not encode any of the *ccm* genes or cyt *c* in its genome. To confirm that *M. barkeri* Fusaro does not produce cyt *c* we assayed the cell lysate with the peroxidase-based heme stain for cyt *c* described earlier. Using this technique, did not detect any signal of cyt *c* in *M. barkeri* Fusaro (Supplementary Figure 7). We built a plasmid-based expression system containing some or all of the following components: i) *ccmABCEF1F2* genes under the control of a tetracycline inducible promoter and ii) *mmcA* (a diagnostic cyt *c*) with a C-terminal TAP tag under the control of a constitutive promoter (Figure 5a). Each of these plasmids was integrated on the *M. barkeri* Fusaro chromosome at a neutral locus using a ØC31 integrase system described previously [30]. We were able to detect a band corresponding to the TAP-tagged MmcA by immunoblotting with anti-FLAG antibodies and heme staining in *M. barkeri* strains expressing both *mmcA* and *ccm* genes (Figure 5b). These results clearly indicate that the *ccmABCEF1F2* genes in *M. acetivorans,* encode a complete, functional and streamlined version of the System I cytochrome *c* maturation machinery previously characterized in Bacteria.

**Figure 5:**
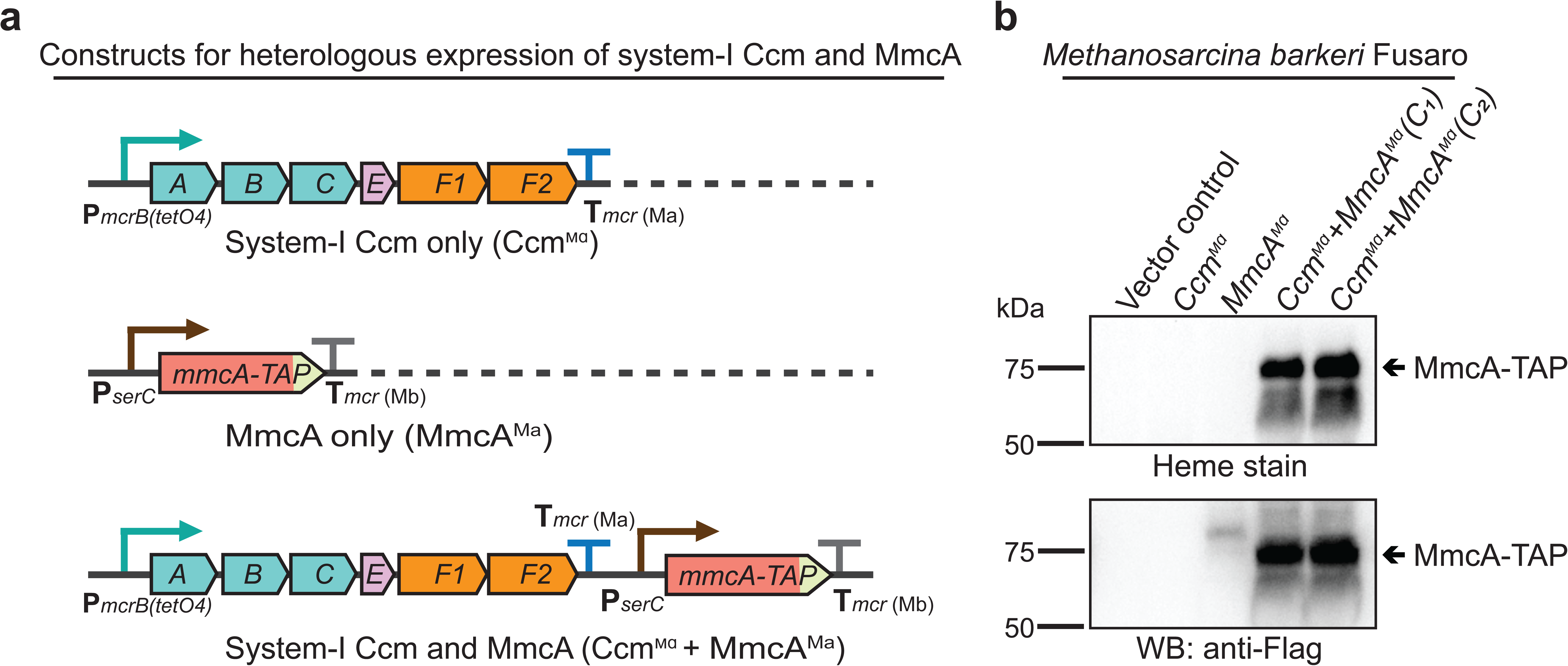
**a)** The design of constructs to express a synthetic operon comprised of the *ccmABCEF1F2* genes from *M. acetivorans* and/or the *mmcA* coding sequence from *M. acetivorans* with a C-terminal tandem affinity purification (TAP) tag comprised of a 3X FLAG sequence and a Twin-Strep sequence. In these constructs, the expression of the *ccmABCEF1F2* operon is driven by a tetracycline-inducible medium-strength promoter, P*mcrB*(*tetO4*), and a transcriptional terminator of the *mcr* operon from *M. acetivorans* T*mcr*(Ma) is provided after the last coding sequence. Each gene, apart from *ccmA,* contains its native Shine-Dalgarno sequence. The C-terminal TAP-tagged MmcA is placed under the control of a constitutive medium-strength promoter (P*serC*) and the transcriptional terminator of the *mcr* operon from *M. barkeri* Tmcr(Mb) has been added immediately downstream of the coding sequence. These constructs are integrated at the ØC31 attachment site at a neutral locus on the *M. barkeri* Fusaro. **b)** Heme peroxidase assays to detect the production of heme-bound Holo-MmcA (top) and Western blots (WB) to detect C-terminal TAP- Tagged MmcA protein (bottom) in whole cell lysates of *M. barkeri* Fusaro cultures grown in medium containing 100 µg/ml tetracycline. A diagnostic *c*-type cytochrome like MmcA can be produced in a heterologous host lacking a native *c*-type cytochrome maturation machinery (*M. barkeri)* by expressing the *ccmABCEF1F2* genes from *M. acetivorans* (Lanes 4-5). C1 and C2 refer to two independently colonies of the *M. barkeri* Fusaro cointegrate expressing the Ccm machinery and MmcA from *M. acetivorans.* An equal amount of whole cell lysate (∼60 µg) was loaded in each lane. The vector control used for this experiment is pJK029A as described elsewhere (Guss *et al*., 2008).

### The production of *c*-type cytochromes is important for growth of *M. acetivorans*

Multiple cyt *c* are encoded in the genome of *M. acetivorans* and some of them, like MmcA, are associated with the electron transport chain; yet the cyt *c* maturation machinery does not seem to be essential under standard growth conditions (i.e. in medium with trimethylamine hydrochloride as the sole carbon and energy source). To determine the physiological role of the Ccm machinery as well as the cyt(s) *c* produced by this process, we assayed the growth characteristics of the Δ*mmcA*, Δ*ccmE* and Δ*ccmABC* mutants on growth substrates that represent a variety of methanogenesis pathways and thermodynamic regimes for *M. acetivorans* (Figure 6 and Supplementary Figure 8). In these analyses, the Δ*ccmE* and Δ*ccmABC* mutants represent lesions in the Ccm pathway that would prevent maturation of all expressed cyt *c,* whereas the Δ*mmcA* mutant represents an in-frame deletion in a specific cyt *c* found in the membrane-associated Rnf (**<u>R</u>**hodobacter **<u>n</u>**itrogen **<u>f</u>**ixation) complex that catalyzes the transfer of electrons between ferredoxin and methanophenazine (a membrane-bound electron carrier) coupled to Na^+^ translocation [26, 34] (Supplementary Figure 2b). We assayed growth on substrates that represent the two modes of methanogenesis in *M. acetivorans:* the methylotrophic pathway and the acetoclastic pathway. During methylotrophic growth, on compounds like trimethylamine hydrochloride (TMA), methanol or dimethyl sulfide (DMS), cells disproportionate the methylated compound to produce carbon dioxide and methane in a 3:1 ratio (Supplementary Figures 8a, 8b and 8d). In contrast, during growth on acetate, via the acetoclastic pathway, the substrate undergoes dismutation to produce carbon dioxide and methane in a 1:1 ratio (Supplementary Figure 8c). Furthermore, the free energy yields (ΔG°’) of these substrates, from -164.5 kJ/mol TMA to -36.0 kJ/mol acetate, also captures the entire bioenergetic landscape of methanogenesis.

**Figure 6:**
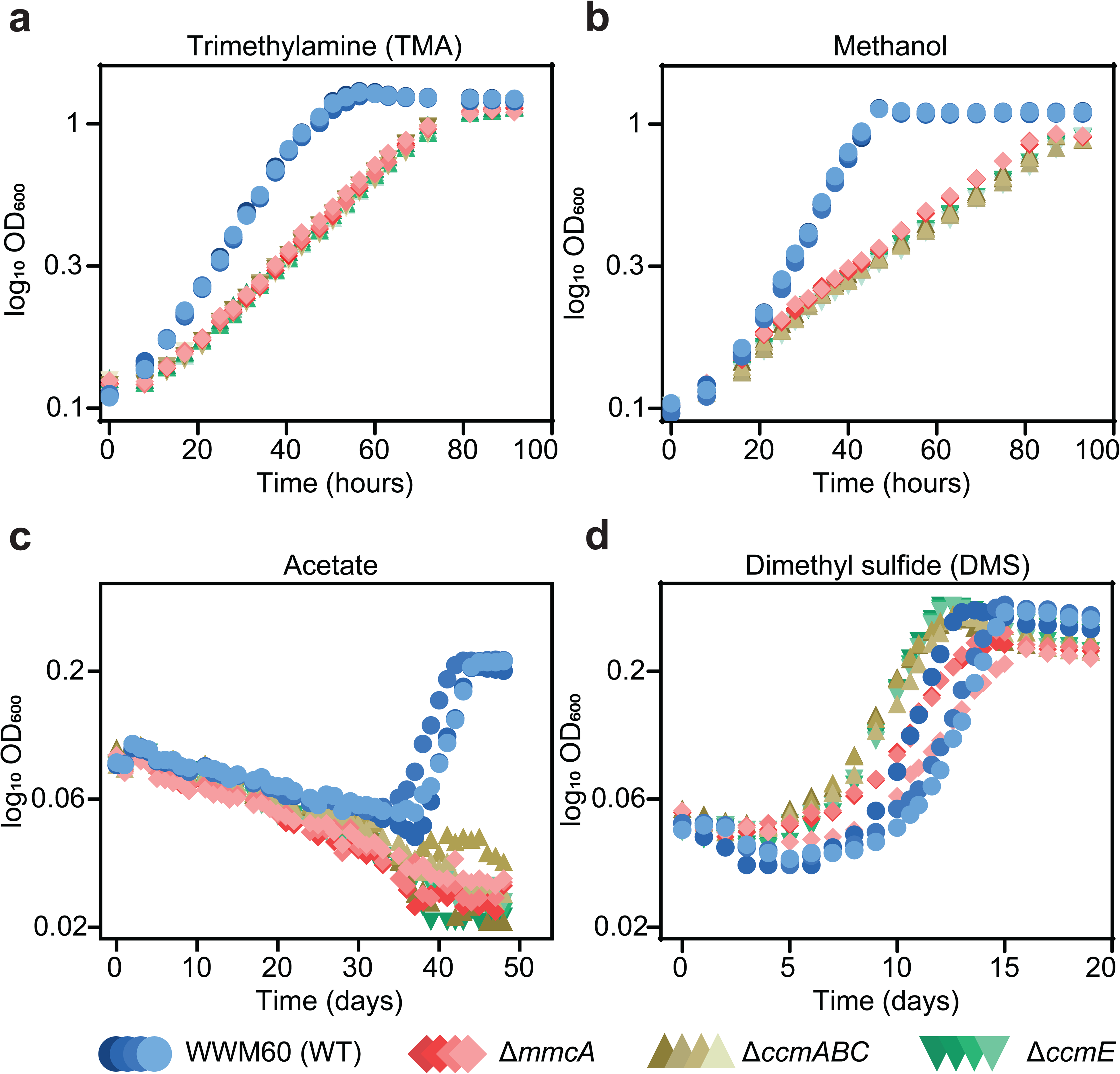
Growth curves of the parent strain (WWM60; referred to as WT) (in blue circles), the Δ*mmcA* mutant (in red diamonds), the Δ*ccmABC* mutant (in olive triangles) and the Δ*ccmE* mutant (in green inverted triangles) in High-Salt minimal medium with **a)** 50 mM trimethylamine hydrochloride (TMA), **b)** 125 mM methanol, **c)** 40 mM sodium acetate (Acetate), and **d)**20 mM dimethylsulfide (DMS) as the sole carbon and energy source. Four replicates were used for growth assays on TMA and methanol and three replicates were used for growth assays on Acetate and DMS.

We observed substantially slower growth than the parent strain for the *ccm*-deletion mutants (Δ*ccmABC* and Δ*ccmE)* as well as the cyt *c* deletion mutant (Δ*mmcA*) under all conditions tested; however, the magnitude of the growth defect varied dramatically (Figure 6; Supplementary Table 2). In growth medium with acetate as the sole substrate, we did not detect any growth for the *ccm*-deletion mutants or the Δ*mmcA* mutant (over the course of 7 weeks), which indicate that holo-MmcA and, possibly, other cyt *c* produced by the Ccm pathway are essential for acetoclastic methanogenesis (Figure 6c; Supplementary Table 2). In contrast, cyt *c* are not essential but important during growth on methylated compounds however, the growth characteristics of each mutant varied substantially (Figures 6a, 6b, and 6d; Supplementary Table 2). Notably, despite a shorter lag, the *ccm*-deletion mutants grew slower than the parent strain on DMS (Figure 6d). On TMA and DMS, the *ccm*-deletion mutants and Δ*mmcA* mutant had a slower growth rate (by a factor of 0.60 and 0.64, respectively) compared to the parent (WWM60) (Figures 6a and 6d; Supplementary Table 2). But, the growth of the *ccm*-deletion mutants and the Δ*mmcA* mutant was indistinguishable from each other (Supplementary Table 2). Taken together, these data indicate that although MmcA is dispensable, it is the sole physiologically relevant cyt *c* during growth on TMA and DMS. In contrast, on methanol, the *ccm-*deletion mutants grew significantly slower than the parent (WWM60) and the Δ*mmcA* mutant as well (by a factor of 0.34 and 0.92, respectively) (Figure 6b; Supplementary Table 2). These data suggest that MmcA is the predominant cyt *c* synthesized on methanol, and is vital for growth, but other cyt *c* encoded in the genome also play a minor but physiologically relevant role. Our growth analyses underscore the importance of cyt *c* on the physiology of *M. acetivorans* and highlight that the role of individual cyt(s) *c* can vary in a substrate-specific manner.

### Evolutionary Analysis of *c*-type Cytochrome biogenesis in Archaea

Even though cyt *c* are important for optimal growth and methanogenesis in *M. acetivorans,* the Ccm machinery and cyt *c* proteins are absent in the close relative *M. barkeri* Fusaro. This uneven distribution of cyt *c* in strains within the Genus *Methanosarcina* could be due to gene loss in strains like *M. barkeri* Fusaro or gene gain by horizontal gene transfer (HGT) of the Ccm machinery and cyt *c* genes in strains like *M. acetivorans*. To distinguish between the two evolutionary hypotheses, we mapped the distribution of the Ccm machinery on the species tree of strains belonging to the genus *Methanosarcina* that are present in the Genome Taxonomy Database (GTDB) [48]. The species tree is based on core genome alignment of the corresponding strains and was obtained from Annotree [49] (Figure 7a). Either all genes of the Ccm machinery are present in a strain or the whole machinery is completely absent (Figure 7a). The strains that encode the Ccm machinery are not monophyletic i.e. the Ccm machinery is not just present in one single clade of *Methanosarcina* strains. This observation strongly supports the hypothesis that the Ccm machinery was present in the last common ancestor of extant *Methanosarcina* spp. and strains like *M. barkeri* Fusaro have lost these genes (Figure 7a). This hypothesis is corroborated by the synteny of the chromosomal locus surrounding the *ccmABC, ccmE,* and *ccmF1ccmF2* genes across all *Methanosarcina* spp. (Figure 7b). Notably, strains like *M. barkeri* MS and *M. vacuolata* contain a truncated ORF (192 bp and 195 bp, respectively) that has sequence homology to the C-terminus of *ccmE* likely as a scar of a gene loss event (Figures 7a and 7b). To test whether the Ccm machinery was aquired by HGT in the last common ancestor of extant strains within the Genus *Methanosarcina* or is present in other Genera within the Family *Methanosarcinaceae,* we mapped the distribution of the Ccm machinery (i.e. the presence of the *ccmABCEF1F2* genes) across the *Methanosarcinaceae* (Supplementary Figure 9). The Ccm machinery is broadly distributed in methanogens and anaerobic methane oxidizing archaea (ANME) across the *Methanosarcinaceae* suggesting an important role for cyt *c* in methane metabolism across this clade (Supplementary Figure 9). Whilst the Ccm machinery is universally conserved in some Genera, like *Methanococcoides, Methanosalsum,* and *Methanohalobium*, this whole machinery seems to have been lost in several members of other Genera, like *Methanosarcina, Methanolobus, Methanoperedens* and *Methanomethylovorans* (Supplementary Figure 9). Thus, it is possible that loss of the Ccm machinery and cyt *c* proteins occurs frequently as an adaptation to certain environmental conditions.

**Figure 7:**
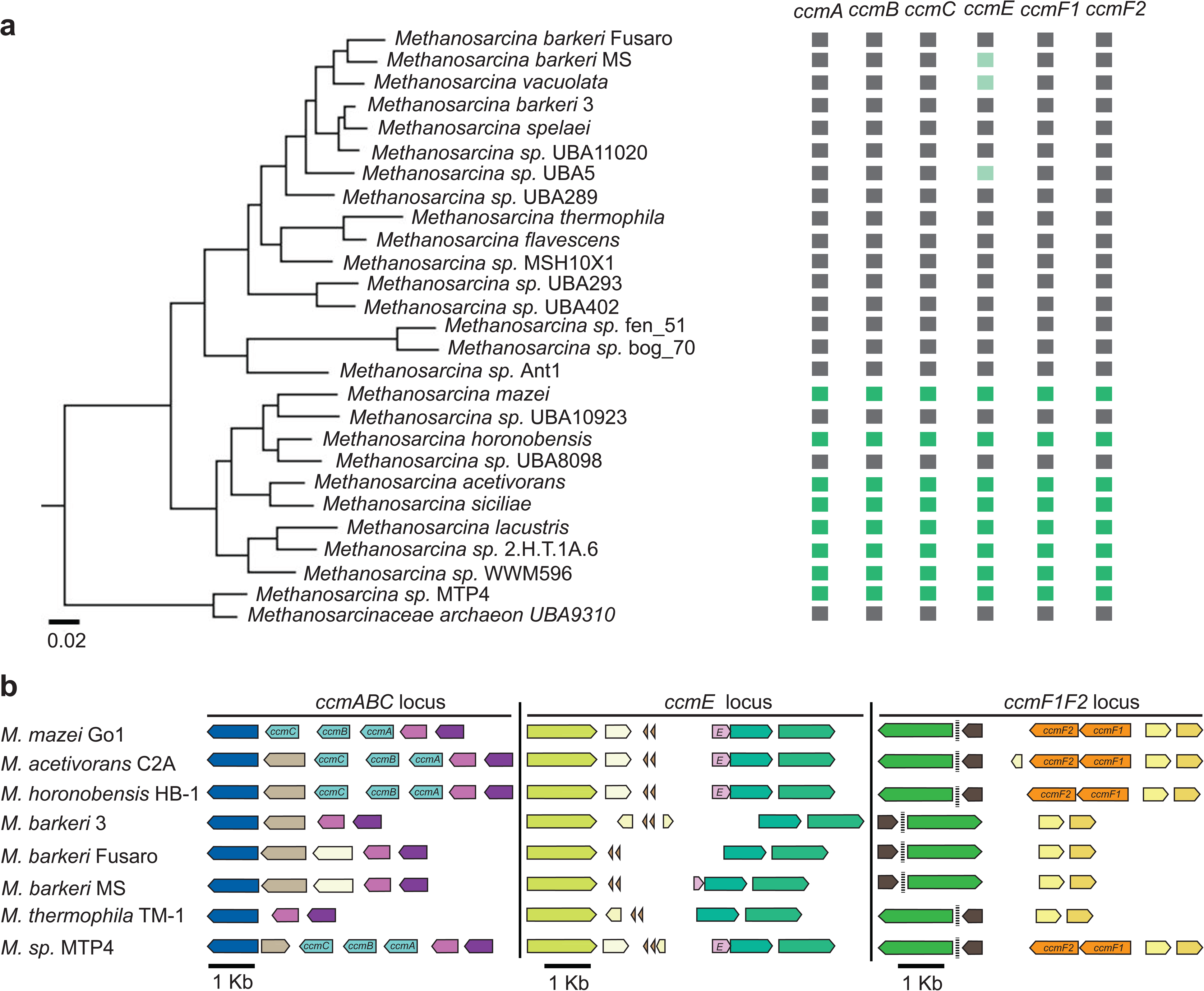
**a)** A phylogenetic tree based on whole genome comparison for strains belonging to the genus *Methanosarcina* in the Genome Taxonomy Database (GTDB) was obtained from AnnoTree (ref). The presence or absence of genes functionally annotated as *ccmA, ccmB, ccmC, ccmE, ccmF1 and ccmF2* in the genome of individual strains is shown in dark green (present) or black (absent) respectively. A few strains like *M. barkeri* MS, *M. vacuolata,* and *M. sp.* UBA5 encode a truncated copy of *ccmE* as indicated in light green. **b)** Chromosomal organization of genes surrounding the *ccmABC, ccmE,* and *ccmF1ccmF2* locus in *Methanosarcina* strains. Genes of the same color (except light yellow) represent members of the same orthologous group. Synteny of the *ccmABC, ccmE,* and *ccmF1ccmF2* chromosomal locus across members of the Genus *Methanosarcina* supports the hypothesis that these genes were present in the last common ancestor and have been lost in several extant lineages.

To test if the Ccm machinery in members of the *Methanosarcinaceae* was vertically inherited from an archaeal ancestor or acquired through inter-domain HGT from bacterial clades, we constructed maximum-likelihood phylogenetic trees of each *ccm* gene from *M. acetivorans* (Figure 8; Supplementary Figures 10-15). For each of the *ccm* genes, one or more clades comprising of strains beloinging to the Order *Methanosarcinales* is nested within clades derived from different bacterial groups (Figure 8; Supplementary Figures 10-15). Based on these trees, it is evident that Ccm machinery in the *Methanosarcinales* was acquired by HGT from bacteria. It is also likely that different *ccm* genes were acquired from different bacteria, which is further corroborated by the observation that these genes are not present in an operon contrary their chromosomal arrangement in bacterial strains (Supplementary Fgure 1, 10-15). Taken together, the streamlined Ccm machinery, which lacks CcmDHI, is not ancestral to the Archaea, rather it has been acquired through multiple inter-domain HGT events between Archaea and Bacteria. While it is not possible to rule out the role of historical contingeny in the notable absence of CcmDHI, it is tempting to speculate that unique features of the archaeal cell render CcmHDIG and CcdA dispensable such that a streamlined Ccm machinery comprising of CcmABCEF is sufficient for cyt *c* maturation in Archaea.

**Figure 8:**
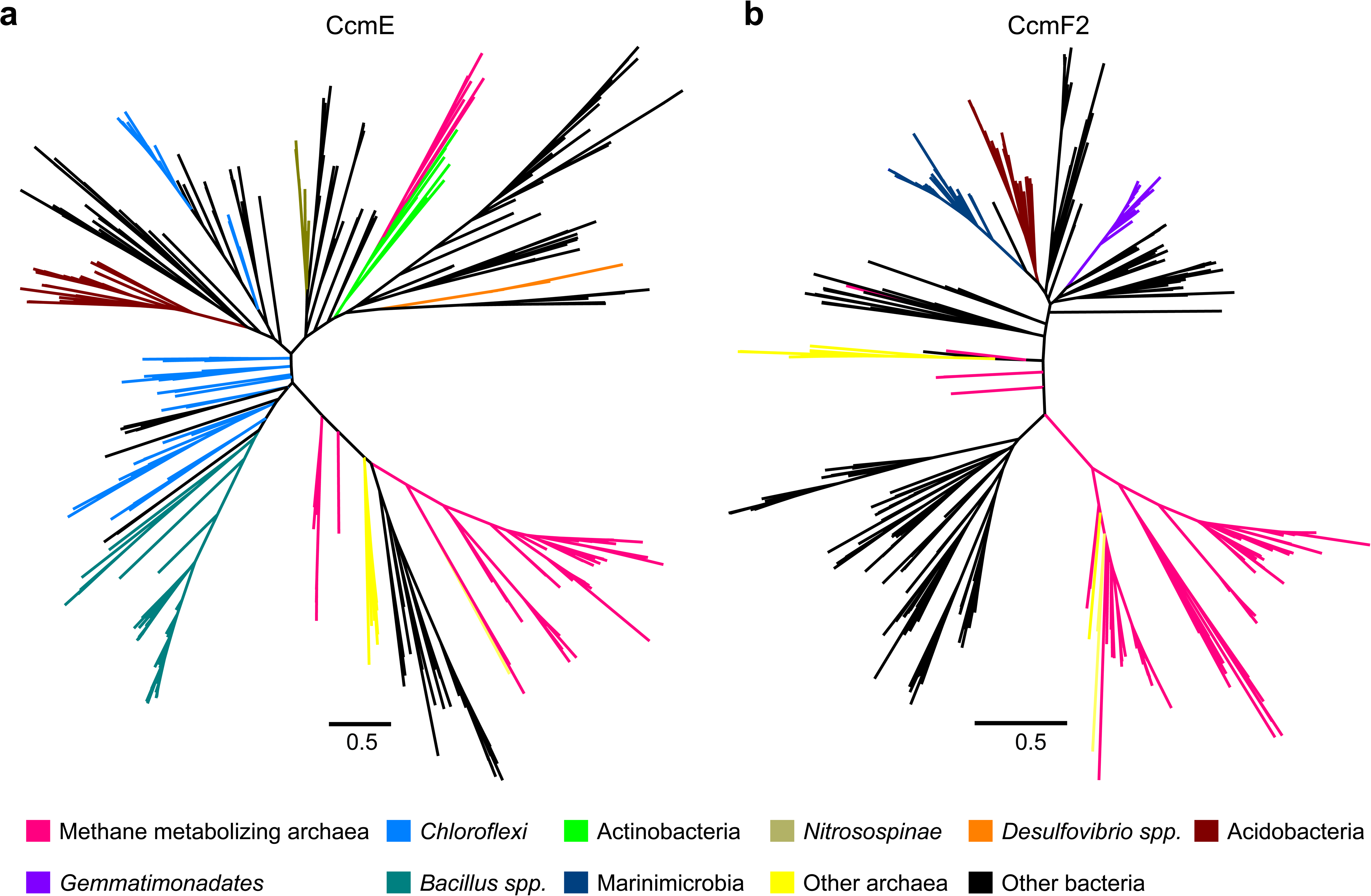
Maximum-likelihood phylogenetic trees of 500 **a)** CcmE and **b)** CcmF2 sequences obtained from the NCBI non-redundant (nr) protein database using the corresponding sequence from *Methanosarcina acetivorans* as the search query. Sequences belonging to certain functional groups or evolutionarily related groups of archaea and bacteria are shown in colors denoted in the legend at the bottom. These gene trees are indicative of multiple Horizontal Gene Transfer (HGT) events between members of the Archaea and Bacteria.

## Discussion

Cyt *c* are widely found in members of the Archaea where they catalyze numerous electron transfer processes that facilitate microbial interactions and fuel biogeochemical cycles. The biogenesis of cyt *c* is complex and is mediated by a dedicated pathway that can vary substantially across the tree of Life. In the vast majority of eukaryotes, a single protein called cyt *c* heme lyase (CCHL) is involved in the biogenesis of cyt *c* [5]. In contrast, bacteria contain more complex pathways that are fine tuned to work with lower concentrations of heme [5]. The most complex of these pathways, called the Ccm machinery, requires at least nine to ten proteins (CcmABCDEFGH(I) and CcdA/DsbD) that work in concert for the biogenesis of cyt *c*. The biochemical details of the Ccm machinery in bacteria has been studied in great detail using model systems over the past few decades and the role of each *ccm* gene is well established at this point [5, 6]. Previous research has shown that archaea encode some components of the bacterial s Ccm machinery [11, 12] however, orthologs of at least three proteins (CcmD, CcmH and CcmI) that are absolutely critical for cyt *c* maturation in bacteria are universally absent in archaeal genomes. Thus, it was unclear how the Ccm machinery is being repurposed (if at all) for cyt *c* biogenesis in archaea. In this study, we use the methanogen *Methanosarcina acetivorans* to characterize a streamlined form of the Ccm system comprised of only five proteins (CcmABCEF) that likely serves as the predominant route for cyt *c* biogenesis in members of the Archaea (Figure 2 and Figure 5).

Based on our findings, there are three key differences between the Ccm machinery in Bacteria and Archaea. While *ccmD* seems to be uniformaly absent in archaeal genomes [11] it is universally present in bacterial genomes and is essential for cyt *c* biogenesis in a number of strains ranging from *E. coli* and *R. capsultus* to *D. desulfuricans* [5, 19, 21]. In bacteria, CcmD is a small (69 aa long) transmembrane protein that associates with CcmABC and facilitates the release of holo-CcmE in the periplasm after heme transport and attachment [21, 46]. In *M. acetivorans,* the CcmABC complex is capable of forming holo-CcmE and releasing it into the pseudo-periplasm in a CcmD independent manner (Figure 3). It is unlikely that the *ccmD* gene has been fused to *ccmA, ccmB,* or *ccmC* based on the sequence length and alignment of the archaeal genes with their bacterial counterparts (Supplementary Figures 10-12). It is also unlikely that the CcmD independent mechanism for the release of holo-CcmE from CcmABC might be linked to the CXXXY heme-binding motif in CcmE because some bacteria, like *D. desulfuricans,* that encode *ccmD* and require this gene product for cyt *c* biogenesis also contain a CXXXY heme-binding motif in CcmE (Figure 4) [19]. One plausible hypothesis for the absence of CcmD might be the unusual architecture of archaeal membranes. The archaeal cytoplasmic membrane is composed of isoprenoid lipids (phytanyl units), compared to phospholipids in bacterial cytoplasmic membranes. This lipid composition might alter how CcmABC and CcmE are anchored into the membrane and render CcmD inconsequential during the formation and release of holo-CcmE.

Next, the archaeal CcmE universally contains a CXXXY heme-binding motif whereas bacterial CcmE can either have a HXXXY motif or a CXXXY motif depending on the strain [5, 19]. In *E. coli,* a H130C allele of CcmE can still bind heme but cannot transfer the heme to the CcmFH complex [19, 50, 51]. In contrast, a C120H mutation in *M. acetivorans* seems to destabilize the protein to such an extent that it can no longer be detected (Figure 4 and Supplementary Figure 5). Similarly, no heme-bound CcmE was detected for the C127H allele of CcmE derived from the bacterium *D. desulfuricans* [19]. Taken together, these data suggest that the Cys and His residues involved in heme-binding CcmE are not interchangeable. In fact, many other as yet uncharacterized features of CcmE, possibly unique to these two groups of CcmE proteins, are likely to be equally important for the formation of holo-CcmE and the transfer of heme to the apo cyt *c* via interactions between CcmE and CcmF as well other proteins. Whether the absence of a HXXXY motif containing CcmE in archaea is just an artifact of historical contingency or due to functional/evolutionary constraints remains unclear.

Finally, the cyt *c* synthetase complex in bacteria contains two (CcmFH or CcmFI) or three (CcmFHI) proteins depending on the strain whereas the archaeal counterpart only contains CcmF. In *M. acetivorans*, the CcmF gene is split into two coding sequences (*ccmF1* and *ccmF2*), each of which might be derived from different bacterial groups (Figure 8 and Supplementary Figures 14, 15). Functionally important domains of the cyt *c* synthetase complex including the two histidine residues that coordiante the pseudo-periplasmic heme *b* as well as the highly conserved WWD domain are all present in CcmF2 (Supplementary Figure 3). CcmF1 probably plays a secondary role in cyt *c* biogenesis, which might explain why a small amount of holo cyt *c* can still be detected in the Δ*ccmF1* mutant of *M. acetivorans* (Figure 2). In bacteria and *Arabidopsis*, CcmH/I have been shown to interact with the apo cyt *c* and deliver it to CcmF [5, 16, 52]. It is unlikely that *M. acetivorans* and other archaea have replaced CcmH/I with a non-orthologous protein as heterlogous expression of *ccmABCEF1F2* in the non-native host, *M. barkeri* Fusaro, seems to be sufficient for cyt *c* biogenesis (Figure 5). However, at present, we cannot distinguish between the possibility that *M. barkeri* Fusaro also encodes this non-ortholgous protein or that the standalone CcmF in *M. acetivorans* performs the function of CcmFH in *E. coli*.

Within the Archaea, cyt *c* are especially enriched in the Family *Methanosarcinaceae* wherein these proteins have likely fostered tremendous metabolic innovation [53]. Our work and other studies [28, 35, 36, 39] show that cyt *c* like MmcA are intricately coupled to growth under a variety of conditions in the marine methanogen *M. acetivorans* (Figure 6). Futhermore, membrane-spanning multi-heme cyt *c* have been shown to faclitate anaerobic methane oxidation by microbial consortia comprised of anaerobic methanotrophic archaea (ANME) and sulfate reducing bacteria (SRB) [54–57] and recent studies also suggest an important role for cyt *c* in faciliating anaerobic alkane metabolism in several members of the *Methanosarcinaceae* [40]. Since ANME and alkane oxidizing archaea are yet to be isolated in pure culture, our ability to maipulate the biogenesis of cyt *c* also renders *M. acetivorans* as an ideal platform to perform functional analyses of diverse cyt *c* across the *Methanosarcinaceae.* Curiously, even though cyt *c* are widely present in members of the *Methanosarcinaceae,* there are many extant strains, such as *M. barkeri* Fusaro, that lack the Ccm machinery altogether and do not encode any cyt *c* proteins in their genome. Our evolutionary analyses provide strong evidence that the uneven distribution of the Ccm machinery and cyt *c* in the *Methanosarcinaceae* is due to gene loss events rather than independent gene gain events (Figure 7; Supplementary Figure 9). It is also worth noting that many of the extant lineages that lack the Ccm machinery and cyt *c* are found in freshwater or anthropogenic sources (like waste digestors); reciprocally, the Ccm machinery and diverse cyt *c* are especially conserved in lineages derived from marine environments (Figure 7; Supplementary Figure 9). This pattern is indicative of an environment-specific selective pressure to retain or discard cyt *c* and the associated biogenesis machinery. However, the specific biotic or abiotic environmental cues that underlie this pattern remain unclear at the moment.

Previous studies [11, 12] have used some of the unique features that unify the archaeal Ccm machinery, notably the absence of CcmDHI and a conserved CXXXY heme binding domain in CcmE, to support an ancestral origin of the Ccm machinery in the Archaea. Our phylogenetic analyses (Figure 8 and Supplementary Figures 10-15) contradict this hypothesis and provide strong evidence supporting the hypothesis that there have been many independent HGT events that have cross-pollinated the Ccm machinery between the Archaea and Bacteria. In fact, even the Ccm machinery in one single archaeon, like *M. acetivorans,* may not have been derived from a single source (Figure 8 and Supplementary Figures 10-15). Taken together, the streamlined Ccm machinery in the Archaea is functionally uniform despite independent evolutionary origins.

## Materials and Methods

### In silico design of single guide RNAs (sgRNAs) for Cas9-mediated genome editing

Twenty bp target sequences for Cas9-mediated genome editing in this study are listed in Supplementary Table 3. All target sequences were chosen using the CRISPR site finder tool in Geneious Prime version 11.0 with the following parameters: *i)* the PAM site was set to NGG at the 3’ end, *ii)* 0 mismatches were allowed against off-targets, and *iii)* 0 mismatches were allowed to be indels. Activity scores were predicted using the methods described previously [58]. The *M. acetivorans* chromosome and the plasmid pC2A were used to score off-target binding sites.

### Plasmid construction

All plasmids used in this study are listed in Supplementary Table 4 and the primers used to generate the plasmids are listed in Supplementary Table 5. Plasmids for Cas9 mediated genome editing were designed using pDN201 as the backbone described previously [32]. Briefly, PCR fragments with the P*mtaCB1* promoter from *M. acetivorans*, sgRNA(s) targeting gene(s) of interest, and the *mtaCB1* terminator from *M. acetivorans* were fused to *AscI* linearized pDN201 using the Gibson assembly method as described previously [32]. Subsequently, PCR fragments with the repair template were fused to the sgRNA containing vector linearized with *PmeI* using the Gibson assembly method as described previously [32]. All plasmid-based overexpression constructs were constructed using pJK029A[30] as the backbone. Fragments containing the gene of interest and promoter, terminators or tandem affinity purification tags as indicated in Supplementary Table 4 were amplified and fused to the P*mcrB*(*tetO4*) promoter in pJK029A [30] linearized with *NdeI* and *HindIII* using the Gibson assembly method as described previously [32]. A cointegrate of the pDN201-derived plasmid or pJK029A-derived plasmid and pAMG40, containing the pC2A origin of replication, was obtained using the Gateway BP Clonase II Enzyme Mix (ThermoFisher Scientific, Waltham, MA, USA) prior to transformation in *Methanosarcina spp.* Standard techniques were used for the isolation and manipulation of plasmid DNA. WM4489, a DH10ß derivative engineered to control copy-number of oriV-based plasmids [59], was used as the host strain for all plasmids generated in this study (Supplementary Table 4). WM4489 was transformed by electroporation at 1.8 kV using an *E. coli* Gene Pulser (Bio-Rad, Hercules, CA). All pDN201-derived plasmids were verified by Sanger sequencing at the UC Berkeley DNA Sequencing Facility and all pAMG40 cointegrates were verified by restriction endonuclease analysis.

### Strains and growth media

All strained used in this study are listed in Supplementary Table 6. All archaeal strains were grown in single-cell morphology [29] at 37°C without shaking in bicarbonate-buffered high-salt (HS) liquid medium with N2/CO2 (80/20) in the headspace. For transformations and growth analyses, 10 mL cultures were grown in Balch tubes with N2/CO2 (80/20) at 8-10 psi in the headspace. For protein purifications, 250 mL cultures were grown in anaerobic bottles with N2/CO2 (80/20) at 3-5 psi in the headspace. All *M. barkeri* Fusaro cells were cultivated in liquid medium supplemented with 125 mM methanol. For mutant generation, *M. acetivorans* and *M. barkeri* Fusaro were plated on agar solidified HS medium (1.6% agar w/v) with 50 mM TMA or 62.5 mM Methanol as the carbon and energy substrate respectively. Solid media plates were incubated in an intra-chamber anaerobic incubator maintained at 37°C with N2/CO2/H2S (79.9/20/0.1) in the headspace, as described previously [60]. Puromycin (RPI, Mount Prospect, IL, USA) to a final concentration of 2 µg/mL and the purine analog 8ADP (CarboSynth, San Diego, CA, USA) to a final concentration of 20 µg/mL were added from sterile, anaerobic stock solutions to select for transformants containing the *pac* (puromycin transacetylase) cassette and to select against the *hpt* (phosphoribosyltransferase) cassette encoded on pC2A-based plasmids, respectively. Anaerobic, sterile stocks of tetracycline hydrochloride in deionized water were prepared fresh shortly before use and added to a final concentration of 100 µg/mL. All mutant strains were verified by Sanger sequencing at the UC Berkeley DNA Sequencing Facility. All *Escherichia coli* strains were grown in Lysogeny broth (LB) or LB-agar at 37 °C with appropriate antibiotics (25 µg/mL kanamycin, and/or 10 µg/mL chloramphenicol) as indicated for different constructs (Supplementary Table 4). All liquid cultures of *E. coli* were grown shaking at 250 rpm. For plasmid preparation, cultures were supplemented with 10 mM rhamnose to increase the plasmid copy number of the pDN201 and pJK029A derived plasmids.

### Transformation of *Methanosarcina spp*

Liposome-mediated transformation was used for *M. acetivorans* and *M. barkeri* Fusaro as described previously [61]. Late-exponential phase culture of *M. acetivorans* (10 mL with TMA) or *M. barkeri* Fusaro (50 mL with MeOH) and 2 µg of plasmid DNA were used for each transformation. Briefly, cells were centrifuged, the supernatant was carefully decanted, and the pellet was resuspended in 1 mL bicarbonate buffered isotonic sucrose (0.85 M) containing 100 µM cysteine. Two µg plasmid DNA and 25 µL DOTAP (N-[1-(2,3- Dioleoyloxy)propyl]-N,N,N-trimethylammonium methylsulfate) (Roche Diagnostics Deutschland GmbH, Mannheim, Germany) were added to the cell suspension and incubated for 4 hours at room temperature in an anaerobic chamber with CO2/H2/N2 (20/4/balance) in the headspace. The mix of cells, plasmid DNA and DOTAP was inoculated in HS-medium with the appropriate growth substrate growth medium and incubated at 37 °C for 12-16 hours prior to plating. *M. acetivorans* cells were spread on puromycin-containing agar-solidified medium using a spreader whereas a *M. barkeri* Fusaro cells were plated used the top-agar method described previously [29, 62].

### Whole genome resequencing of CRISPR-edited *Methanosarcina acetivorans* mutants

A 10 mL culture of DDN029 in HS + 50 mM trimethylamine hydrochloride incubated at 37 °C was harvested at late-exponential phase (OD600 ∼0.8). Genomic DNA was extracted using the Qiagen blood and tissue kit (Qiagen, Hilden, Germany) and the concentration of genomic DNA was measured using a Nanodrop One Microvolume UV-Vis Spectrophotmeter (Thermo Scientific, Waltham, MA, USA). Library preparation and Illumina sequencing was performed at the Microbial Genome Sequencing Center, Pittsburgh, PA, USA. Illumina sequencing reads were aligned to the *Methanosarcina acetivorans* C2A genome and mutations were identified using Breseq version 0.35.5 [63]. Illumina sequencing reads for DDN029 have been deposited to the Sequencing Reads Archive (SRA) with the following BioProject accession number: PRJNA800036.

### Growth assays for *M. acetivorans*

*M. acetivorans* strains were grown in single-cell morphology [29] in bicarbonate-buffered high salt (HS) liquid medium containing 125 mM methanol, 50 mM trimethylamine hydrochloride (TMA), 40 mM sodium acetate, or 20 mM dimethylsulfide (DMS). Most substrates were added to the medium prior to sterilization. DMS was added from an anaerobic stock solution maintained at 4 °C immediately prior to inoculation. For growth analyses, 10 mL cultures were grown in sealed Balch tubes with N2/CO2 (80/20) at 8-10 psi in the headspace. Growth measurements were conducted with at least three independent biological replicates derived from colony-purified isolates using optical density readings at 600 nm (OD600). Optical density readings were obtained from a UV-Vis Spectrophotometer (Gensys 50, Thermo Fisher Scientific, Waltham, MA, USA) outfitted with an adjustable test tube holder that could directly take OD600 readings from cultures in a Balch tube. A Balch tube containing 10mL HS medium with the appropriate growth substrate was used as the ‘Blank’ for OD600 measurements. For growth on methanol, TMA, and DMS, cells were acclimated to the growth substrate for a minimum of five generations prior to quantitative growth measurements. Growth measurements on acetate were performed with cells transferred from HS+TMA medium. A 1:20 dilution of mid-late-exponential phase cultures was used as the inoculum for growth analyses. Growth curves were plotted on a log base 10 scale and growth rate measurements were performed individually for each replicate by calculating the slope of the line that fits as many data points on the growth curve for a linear regression coefficient (R^2^) ≥ 0.99. For maximum OD600 measurements, cells were diluted 1:10 in HS-medium containing the appropriate growth substrate. Growth curve plots and statistical analyses were obtained using GraphPad Prism 9.0.0.

### Purification of membrane fraction containing Tandem Affinity Purification (TAP) tagged CcmE

Protein purification was performed with 250 mL of mid-exponential phase culture grown in HS + 50 mM TMA at 36°C without shaking under aerobic conditions. Cells were harvested by centrifugation (6000 × g) for 20 min at 4 °C and was lysed in 10 mL of Wash buffer (50 mM NaH2PO4, 300 mM NaCl, pH = 8.0) on ice. The cell lysate was clarified by centrifugation at 10,000 × g for 10 min at 4°C, followed by separation of soluble and membrane fractions via high-speed ultracentrifugation at 100,000 × g for 45 min at 4 °C. The membrane pellets were solubilized in 4 mL Wash buffer with 1% Triton 100-X (Sigma-Aldrich, Saint Louis, MO, USA). The solubilized membrane fraction was loaded on a column containing 0.5 mL Strep-tactin Superflow plus resin (50% suspension; Qiagen, Hilden, Germany) equilibrated with 4 mL of the Wash buffer. The column was washed twice with 4 mL of Wash buffer by gravity flow and the purified protein was eluted in four fractions with 0.5 mL Elution buffer (50 mM NaH2PO4, 300 mM NaCl, 2.5 mM desthiobiotin, pH = 8.0) per fraction. The protein was concentrated using 10kDa Amicon filter (Merck Millipore, Burlington, MA, USA). The protein concentration in each fraction was estimated using the Bradford reagent (Sigma-Aldrich, Saint Louis, MO, USA) with BSA (bovine serum albumin) as the standard per the manufacturer’s instructions.

### Heme staining and immunoblotting

Heme staining using heme peroxidase assays were performed as described previously [41, 64]. For heme staining MmcA, total cell lysate of *M. acetivorans* or *M. barkeri* Fusaro was incubated at 65°C for 4 minutes. For heme staining CcmE, cell lysate/enriched-protein samples were not heat treated. Protein samples were resolved by 12% Mini-Protean TGX denaturing SDS-PAGE gel (Bio-Rad, Hercules, CA, USA) and transblotted to 0.2 µm PVDF membrane (Bio-Rad, Hercules, CA, USA) using Trans-Blot Turbo transfer system (Bio-Rad, Hercules, CA, USA). For heme staining, the SuperSignal West Femto kit (Thermo Scientific, Waltham, MA, USA) was used to detect the heme signal on the transblotted PVDF membrane and imaging was performed with ChemiDoc MP Imaging System (Bio-Rad, Hercules, CA, USA). All the heme stain blots were washed with 50 mL stripping buffer (60 mM Tris pH= 7 containing 2% SDS and 7 µL/mL β-mercaptoethanol) shaking at 50 rpm for 1 hour at 50 °C and confirmed for the absence of any peroxidase-based signal from heme before they were used for immunoblots. FLAG-tagged proteins were probed with immunoblotting using monoclonal anti-Flag M2-Peroxidase (HRP) antibody (Sigma-Aldrich, Saint Louis, MO, USA) (1/50,000X dilution) and Immobilon Western Chemiluminescent HRP Substrate (Millipore, Burlington, MA, USA) was used for signal detection. Imaging was performed with ChemiDoc MP Imaging System (Bio-Rad, Hercules, CA, USA). Near equal loading of total protein was estimated using the Bradford reagent (Sigma-Aldrich, Saint Louis, MO, USA) with BSA (bovine serum albumin) as the standard per the manufacturer’s instructions.

### Bioinformatics analyses

Phylogenetic trees of *Methanosarcina* spp. and other strains within the *Methanosarcinales* (as depicted in Figure 7a and Supplementary Figure 9) were obtained from AnnoTree [49]. Annotree was used for functional annotation of *ccm* genes using the following KEGG annotations: *ccmA* (K02913), *ccmB* (K02194), *ccmC* (K02195), *ccmE* (K02197), *ccmF* (02198). For gene trees, 500 closest homologs were extracted from the NCBI non-redundant protein database using the corresponding Ccm gene sequence from *M. acetivorans* as the query in BLAST-P searches. The amino acid sequences of these proteins were aligned using MUSCLE 3.7 on the CIPRES Science Gateway cluster V3.3 [65]. Maximum-likelihood trees were generated using RAxML v8.2.12 with the Jones-Taylor-Thornton (JTT) substitution matrix on the CIPRES Science Gateway cluster V3.3 [65]. Trees were displayed using Fig Tree v1.4.3 (http://tree.bio.ed.ac.uk/software/figtree/). Gene ortholog neighborhood was obtained using the bidirectional best hits for the corresponding *ccm* gene using the Integrated Microbial Genomes and Microbiomes (IMG) platform containing annotated isolate genome and metagenome datasets sequenced at the Joint Genome Institute (JGI)[66]

## Author contributions

D.G. contributed to conceptualization, data curation, formal analysis, methodology, and writing. K.E.S. contributed to data curation, formal analysis, and methodology. D.D.N contributed to conceptualization, data curation, formal analysis, supervision, funding acquisition, project administration, methodology, and writing.

## Acknowledgements

The authors would like to acknowledge members of the Nayak lab for their feedback and input.

## Supplementary Material

**Supplementary Figure 1:**
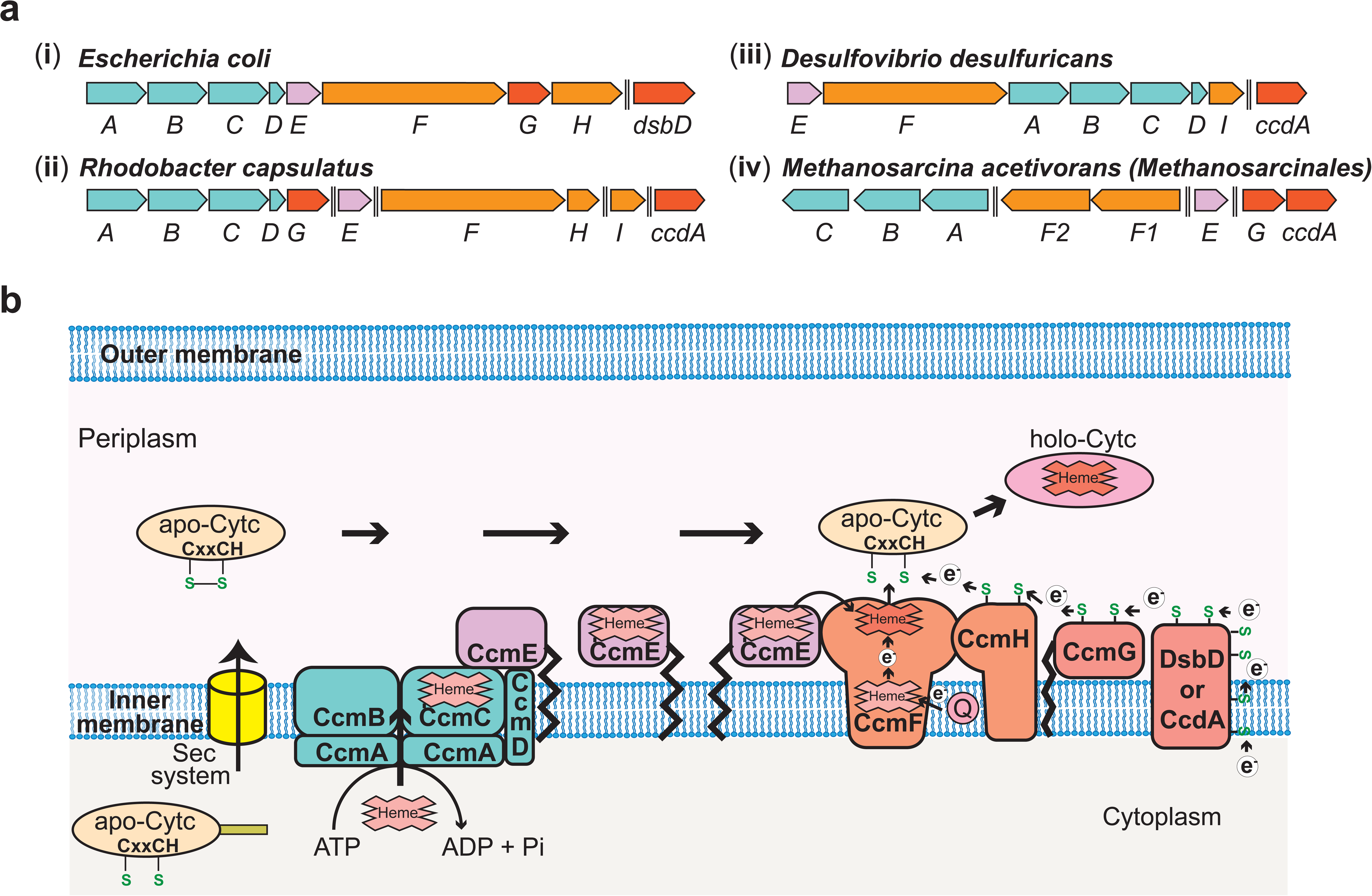
**a)** Chromosomal organization of the System I *c*-type cytochrome maturation (Ccm) machinery genes in a few representative bacteria and the archaeon, *Methanosarcina acetivorans*. In bacteria like **i)** *Escherichia coli* **ii)** *Rhodobacter capsulatus* and **iii)** *Desulfovibrio desulfuricans*, 9-10 genes (*ccmABCDEFGHI* and *dsbD*/*ccdA)* are involved in the maturation of *c*-type cytochromes (cyt *c*), and most of the *ccm* genes are found in an operon. **iv)** The *ccm* genes in the archaeon *M. acetivorans* and other methane metabolizing archaea belonging to the Order *Methanosarcinales* are not found in a single operon and, notably, homologs of three essential genes, *ccmD, ccmH, and ccmI,* are absent. Double dashed lines indicate a distance >110 kbp on the chromosome. **b)** An overview of the System I Ccm machinery from *E. coli.* The apo cyt-*c* is transported across the inner cytoplasmic membrane into the periplasmic space using the Sec system. The CcmABCD membrane-bound complex transports heme *b* to form heme-bound holo-CcmE in an ATP-dependent manner. Holo-CcmE delivers heme *b* to the membrane-bound cyt *c* synthetase complex, CcmFH, that catalyzes the formation of holo cyt *c*. CcmG and DsbD/CcdA facilitate the transfer of electrons to reduce the disulfide bond between the cysteine residues in the CXXCH motif of the apo cyt *c* prior to the formation of holo cyt *c*. Figure adapted from Kranz *et al.*, 2009 (1).

**Supplementary Figure 2:**
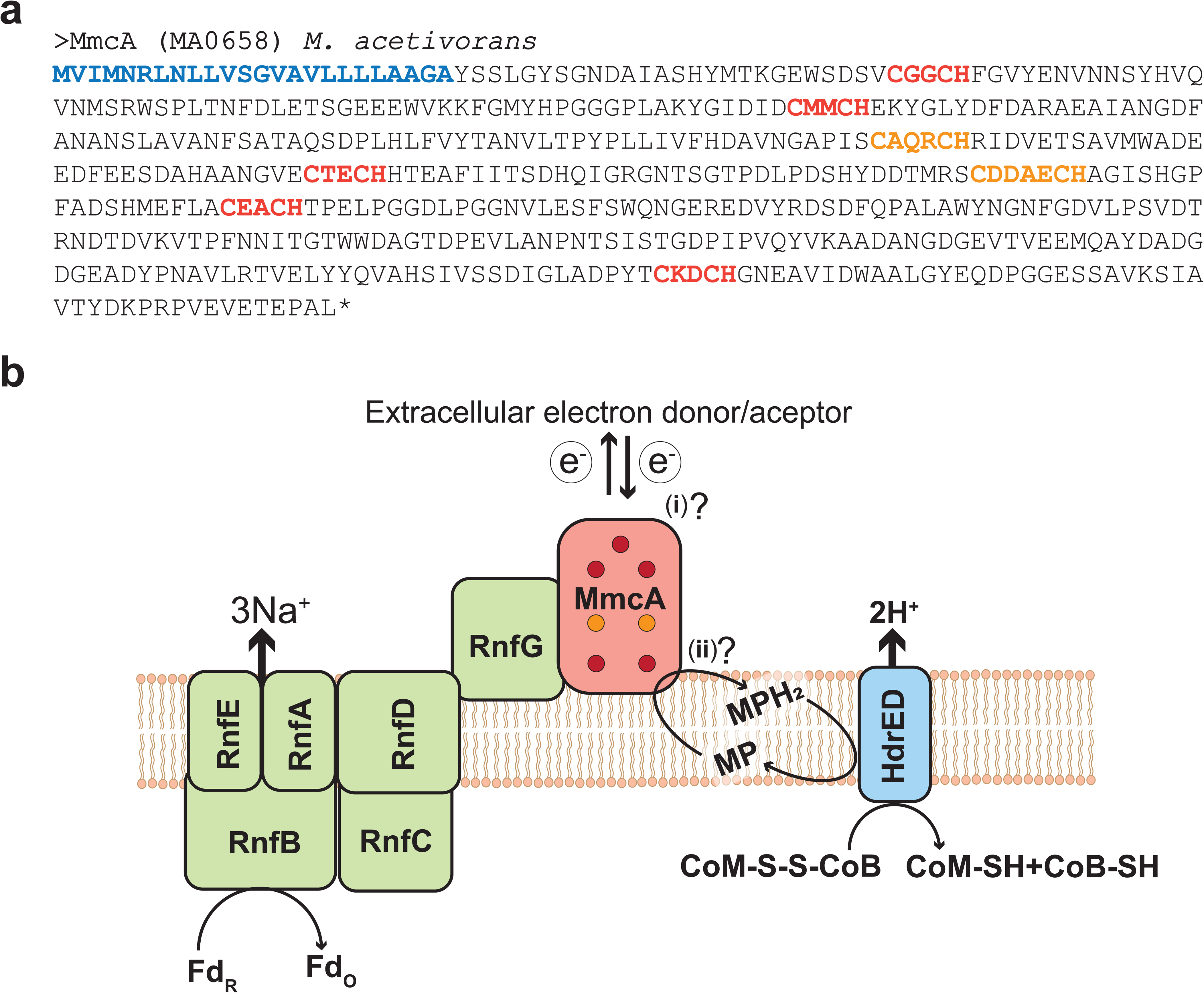
**a)** Amino acid sequence of the *c-*type cytochrome (cyt *c*) MmcA from *Methanosarcina acetivorans.* The predicted signal peptide is shown in blue, five canonical heme-binding motifs (CXXCH) are shown in red and two non-canonical heme-binding motifs (CXXXCH and CXXXXCH) are shown in orange. Based on these observations, MmcA is a heptaheme cyt *c* that is present in the pseudo-periplasm of *M. acetivorans.* **b)** MmcA is associated with the **<u>R</u>**hodobacter **<u>n</u>**itrogen **f**ixation (Rnf) bioenergetic complex in methanogens and other alkane metabolizing archaea. The Rnf complex catalyzes the electron transfer reaction between cytosolic Ferredoxin (Fd) and the membrane-associated electron carrier Methanophenazine (MP) coupled to the translocation of Na^+^ ions across the cytoplasmic membrane. MmcA has been hypothesized to play two roles in the context of the Rnf bioenergetic complex: i) it is involved in the extracellular transfer of electrons from/to Fd or ii) it directly interacts with MP to mediate the final step in the intracellular electron transport reactions from Fd to MP. The reduced MPH2 generated by the Rnf complex is regenerated by the membrane-bound Heterodisulfide reductase (Hdr) that couples the translocation of protons (H^+^) to the transfer of electrons from MP to the CoM-S-S-CoB disulfide that acts as the terminal electron acceptor in cytochrome-containing methanogens like *Methanosarcina acetivorans*.

**Supplementary Figure 3:**
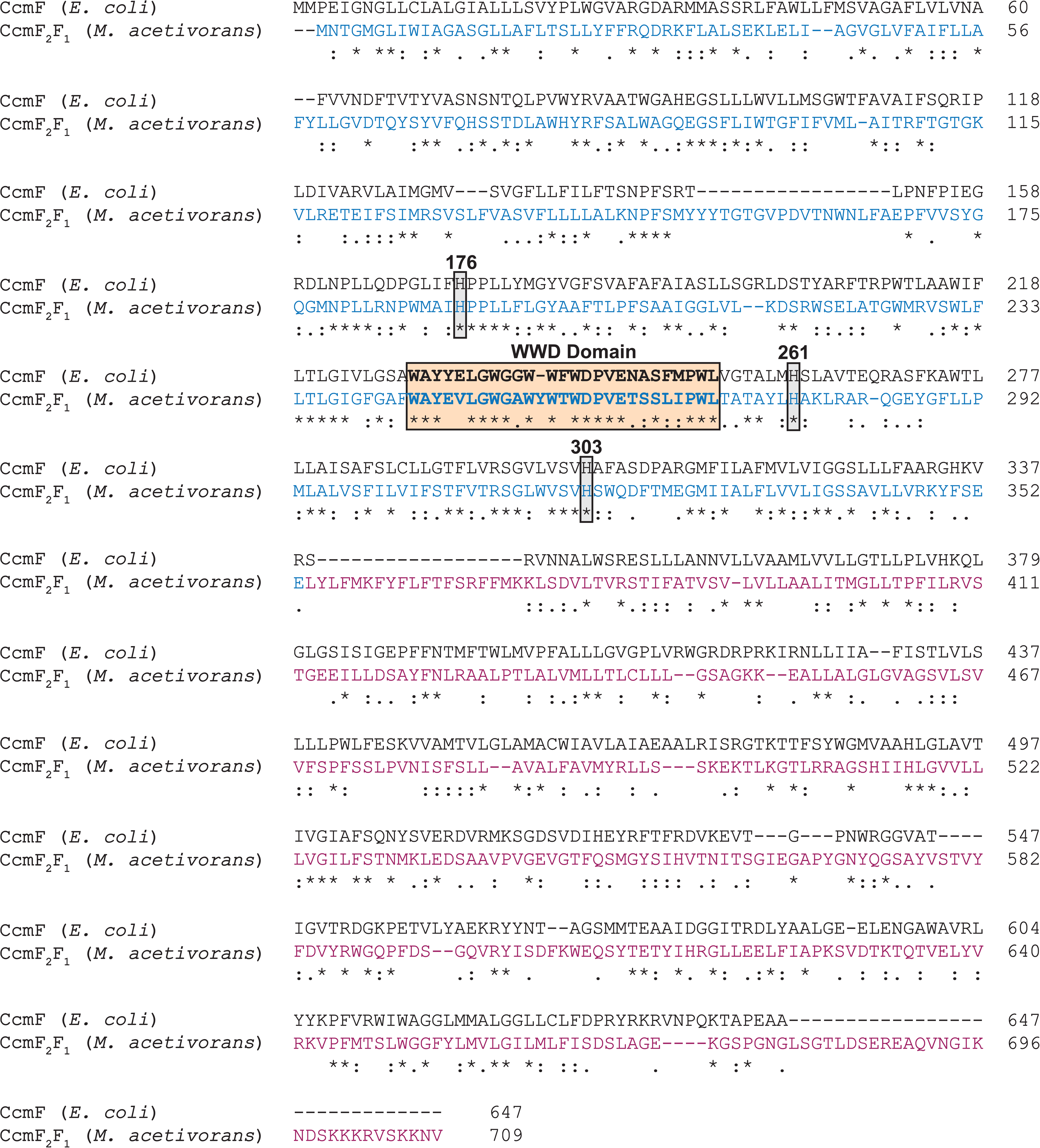
Alignment of the CcmF amino acid sequence from *Escherichia coli* with the concatenated CcmF2-CcmF1 amino acid sequences from *Methanosarcina acetivorans.* Based on this alignment, the CcmF2 sequence (in blue) from *M. acetivorans* is homologous to N-terminus of CcmF from bacteria and the CcmF1 (in purple) sequence of *M. acetivorans* is homologous to the C-terminus of CcmF from bacteria. The highly conserved WWD domain, and three conserved histidine residues that serve as axial ligands for heme *b* in the membrane (His261 in *E. coli*) and the periplasm (His176 and His 303 in *E.* coli) are all conserved in the CcmF2 sequence of *M. acetivorans*.

**Supplementary Figure 4:**
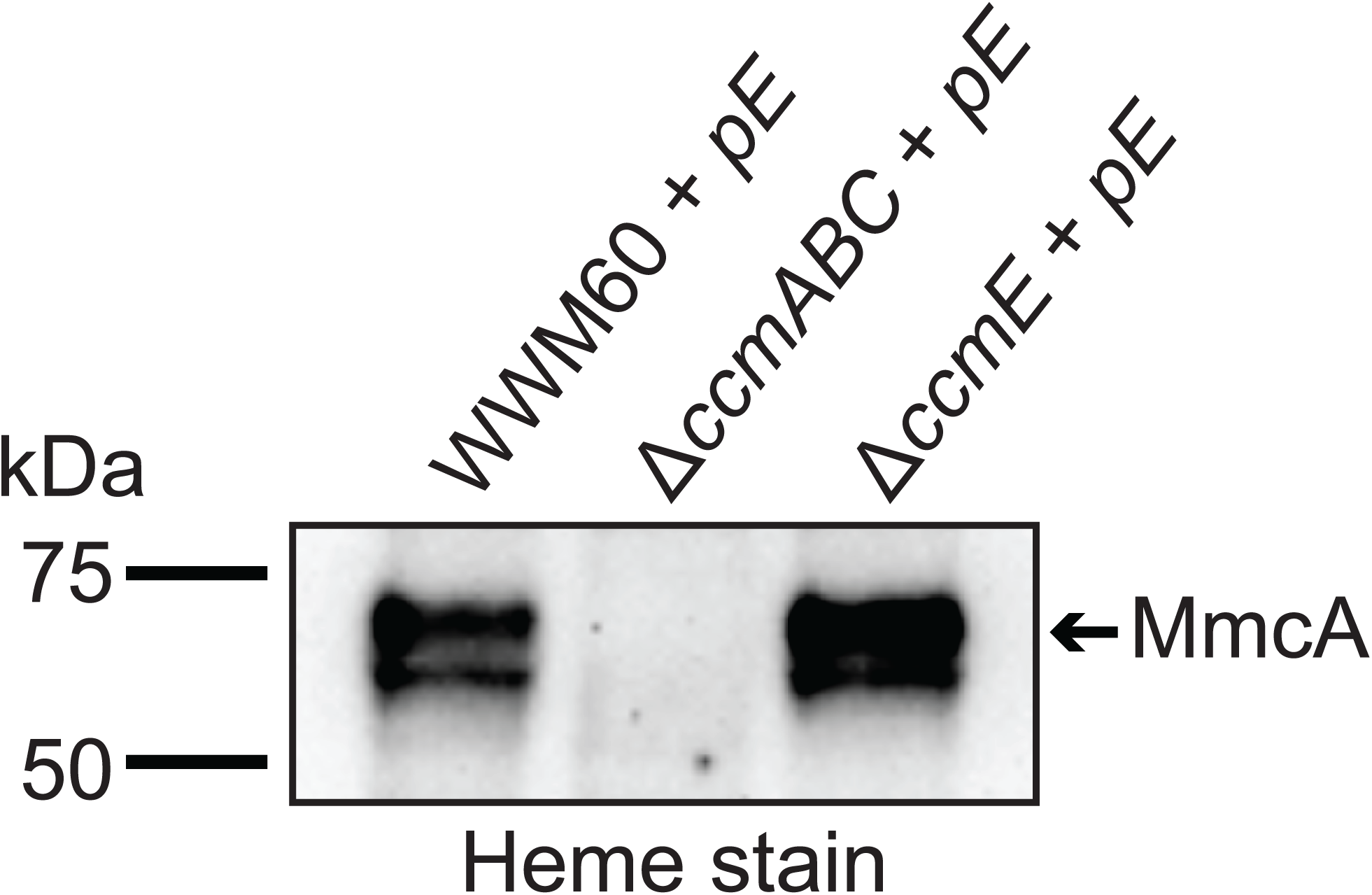
Expression of CcmE with a C-terminal 1XFLAG-1X Strep tag from a plasmid (pE) in the parent strain of *Methanosarcina acetivorans* (WWM60) and the Δ*ccmE* background leads to the production of heme-bound holo-MmcA as detected by a heme peroxidase assay of whole cell lysates (Lanes 1, 3 of heme-stained gel). Thus, the C-tagged CcmE is functional and facilitates the maturation of native *c*-type cytochromes (cyt *c*) like MmcA. In contrast, expression of C-tagged CcmE in the Δ*ccmABC* background does not produce any holo-MmcA (Lane 2 of heme stained gel). In the absence of CcmABC, holo-CcmE does not form, which, in turn, prevents the maturation of native cyt *c* like MmcA. All cultures were grown in medium containing 100 µg/ml tetracycline to induce the expression of C-tagged CcmE from the plasmid pE. An equal amount of protein (∼20 µg) was loaded in each lane.

**Supplementary Figure 5:**
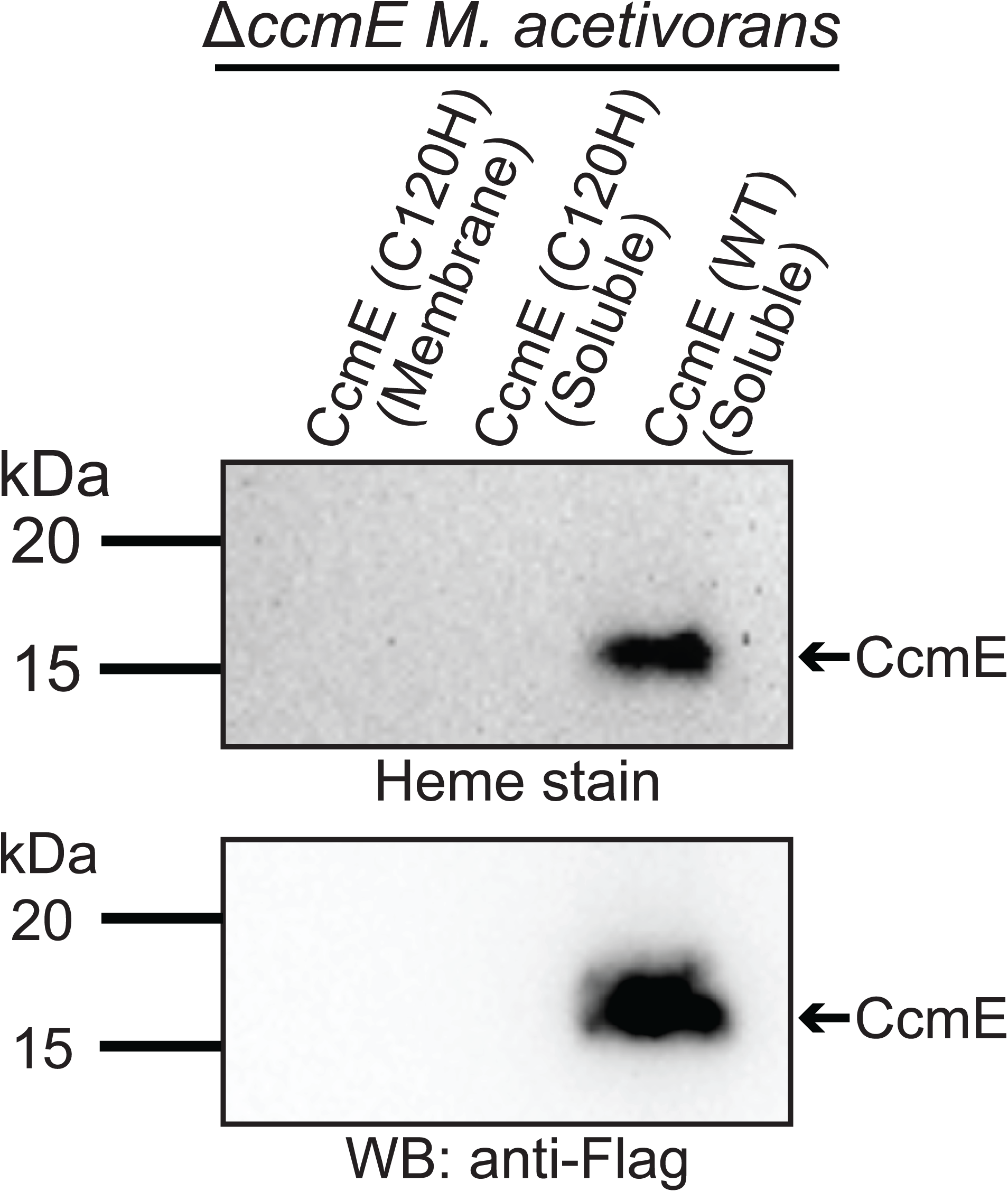
Expression of the C120H mutant of CcmE from *Methanosarcina acetivorans* with a C-terminal 1XFLAG-1X Strep tag in the Δ*ccmE* background does not lead to the production of any CcmE protein that can be detected by Heme peroxidase assays (top gel) or Western Blot using an anti-Flag antibody (bottom gel) in the Strep-enriched membrane fraction (Lane 1) or the soluble fraction (Lane 2). In contrast, expression of the WT sequence CcmE from *M.acetivorans* with a C-terminal 1XFLAG-1X Strep tag in the Δ*ccmE* background leads to the production of protein that can be detected by Heme peroxidase assays (top gel) and Western Blot using an anti-Flag antibody (bottom gel) in the Strep-enriched soluble fraction (Lane 3) and the membrane fraction (see Figure 4). Strep-enriched membrane or soluble fraction represents protein eluted after passing the membrane or the soluble fraction through a Strep-affinity column. The C120H mutant was sequenced verified prior to protein purification. Based on these data, it is highly likely that the C120H substitution destabilizes the CcmE protein. All cultures were grown in medium containing 100 µg/ml tetracycline to induce the expression of the tagged CcmE protein. An equal amount of protein (∼1 µg) was loaded in each lane.

**Supplementary Figure 6:**
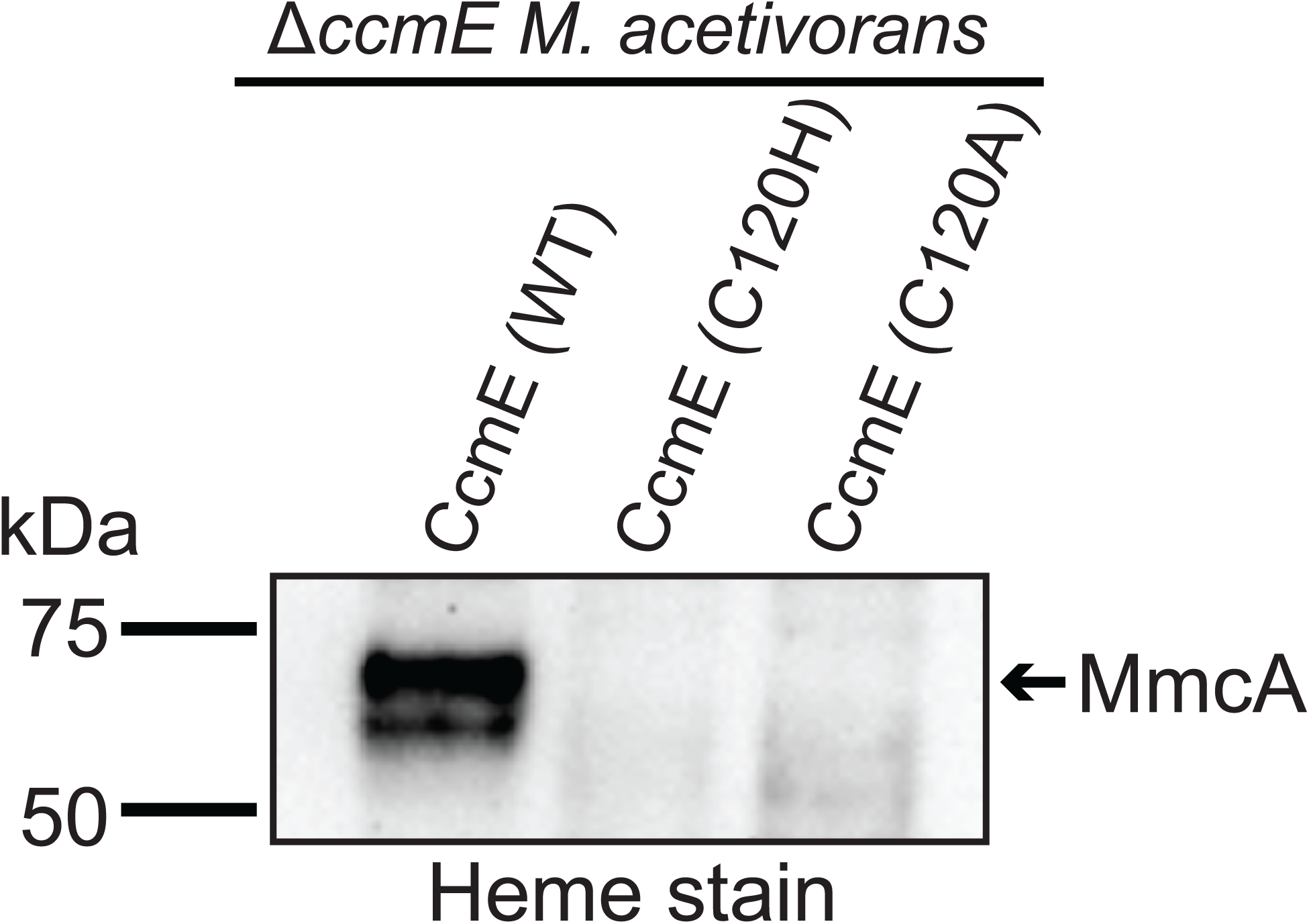
Expression of the wildtype (WT) sequence of CcmE from *Methanosarcina acetivorans* with a C-terminal 1XFLAG-1X Strep tag (C-tagged CcmE) in the Δ*ccmE* background leads to the production of heme-bound holo-MmcA as detected by a heme peroxidase assay of whole cell lysates (Lane 1 of heme-stained gel). Thus, the C-tagged WT sequence of CcmE is functional and facilitates the maturation of native *c*-type cytochromes (cyt *c*) like MmcA. In contrast, expression of C-tagged C120H or C120A mutants of CcmE in the Δ*ccmE* background does not produce any holo-MmcA (Lanes 2,3 of heme stained gel). Thus, the C120H and C120A mutants disrupt the function of CcmE, which, in turn, prevents the maturation of native cyt *c* like MmcA. All cultures were grown in medium containing 100 µg/ml tetracycline to induce the expression of C-tagged CcmE. An equal amount of protein (∼20 µg) was loaded in each lane.

**Supplementary Figure 7:**
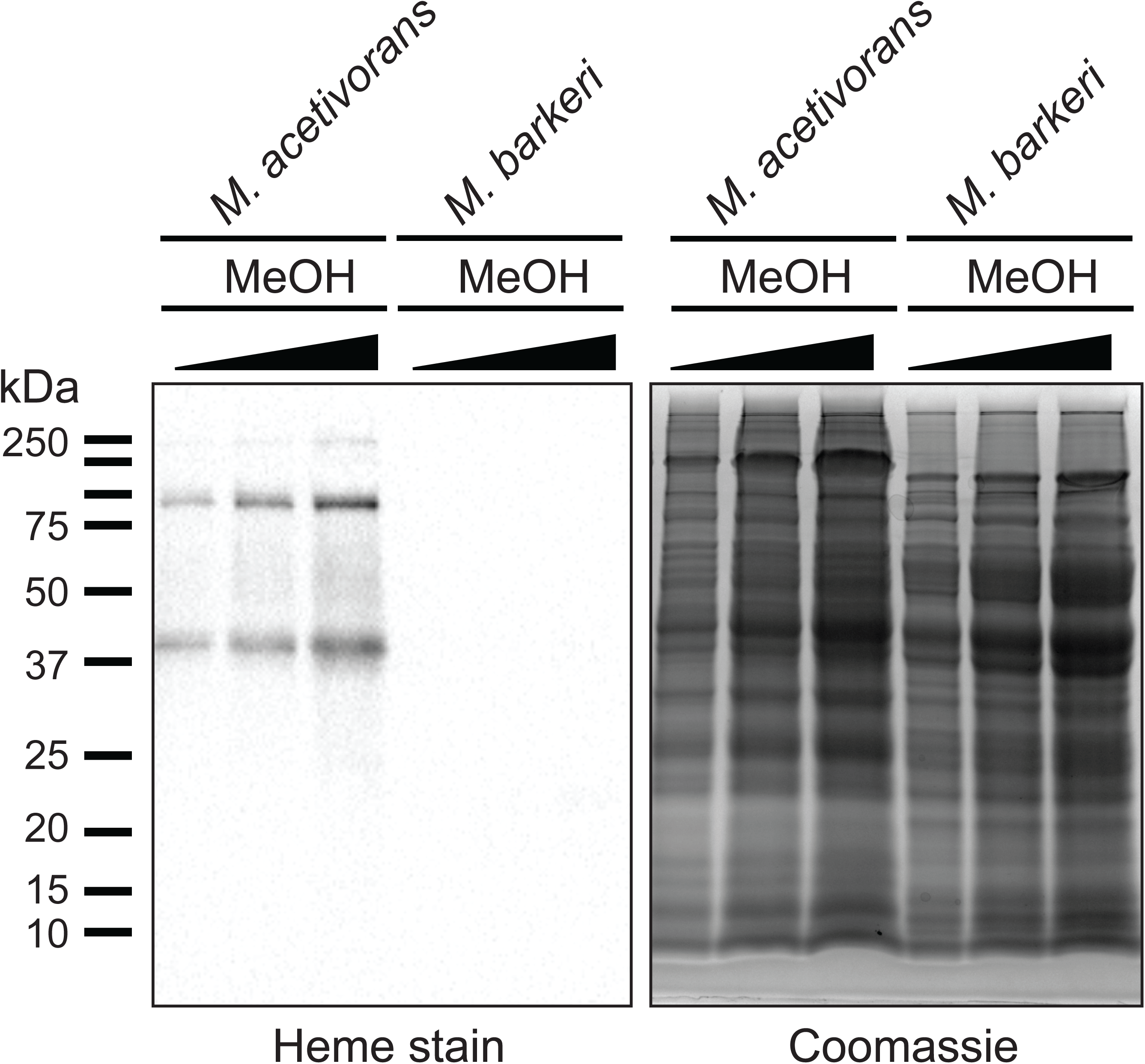
(Left) Heme-peroxidase assays to detect proteins that covalent bind heme (including *c*-type cytochromes) in whole cell lysates of *Methanosarcina acetivorans* (Lanes 1, 2, 3) and *Methanosarcina barkeri* (Lanes 4, 5, 6) grown in high-salt minimal medium with 125 mM methanol as the sole source of carbon and energy. Three different concentrations (∼30, 60, 90 µg) of whole cell lysate were used per strain for these assays as shown in the Coomassie gel control (right).

**Supplementary Figure 8:**
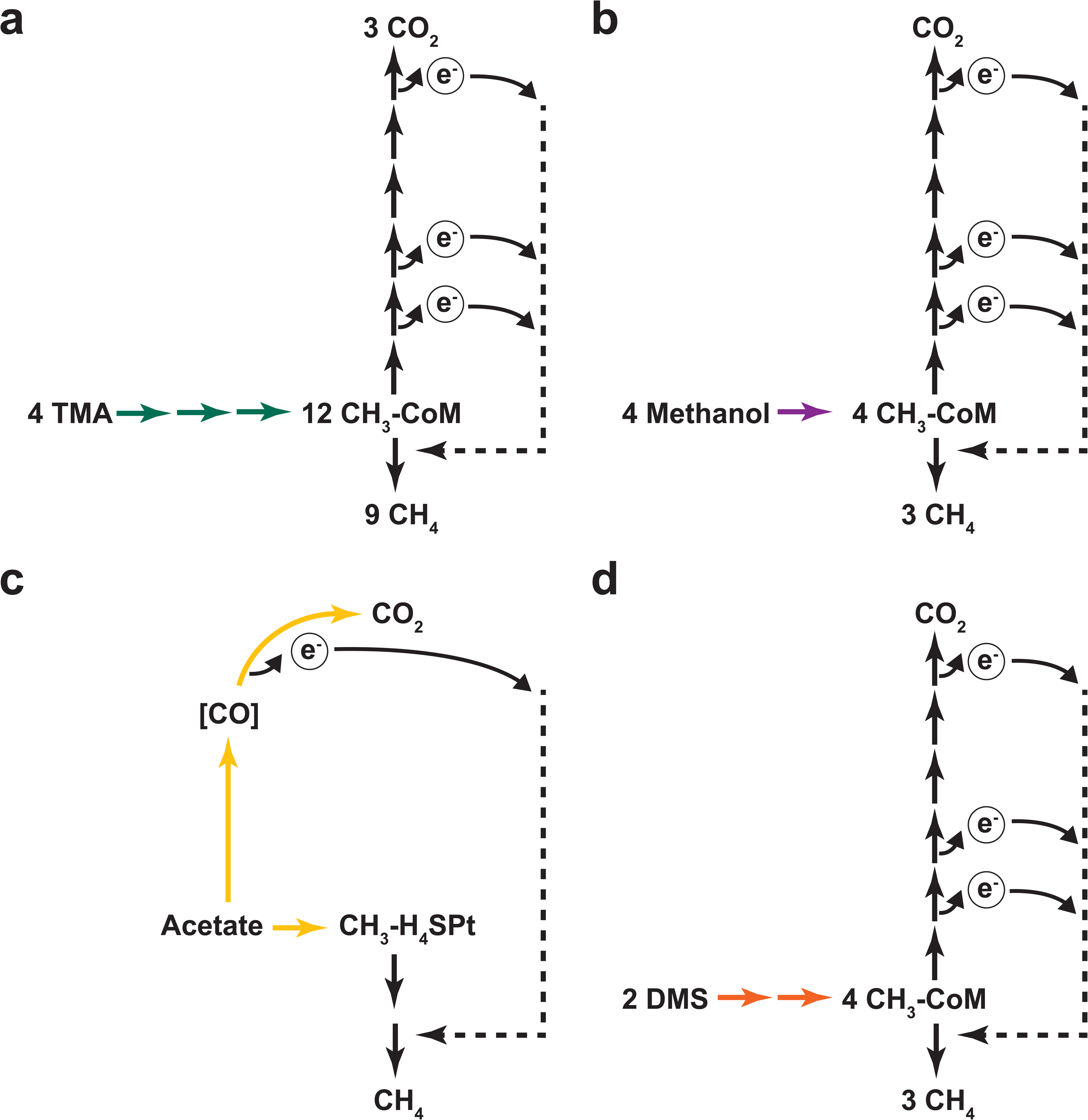
**a)** Methanogenic metabolism of trimethylamine (TMA) in *Methanosarcina acetivorans*. Green arrows represent TMA, dimethylamine (DMA), monomethylamine (MMA) specific methyltransferase reactions that catalyze the transfer of each of the methyl groups in TMA to ultimately form methyl-CoM (CH3-CoM). CH3-CoM undergoes a disproportionation reaction to generate CO2 and CH4 in a 3:1 ratio. **b)** Methanogenic metabolism of methanol in *Methanosarcina acetivorans*. The purple arrow represents the methanol specific reaction that catalyzes the transfer of the methyl group to ultimately form methyl-CoM (CH3-CoM). CH3-CoM undergoes a disproportionation reaction to generate CO2 and CH4 in a 3:1 ratio. **c)** Methanogenic metabolism of acetate in *Methanosarcina acetivorans*. The yellow arrow represents the reaction catalyzed by the acetyl-CoA decarbonylase/synthase (ACDS) enzyme complex that splits acetate into an enzyme bound carbonyl group and a methyl group that is transferred to the C1 carrier tetrahydrosarcinopterin (H4SPt). Oxidation of the carbonyl group to CO2 produces electrons that can be used to reduce the methyl group to CH^4^. During acetoclastic methanogenesis, CO2 and CH4 are produced in a 1:1 ratio. **d)** Methanogenic metabolism of dimethyl sulfide (DMS) in *Methanosarcina acetivorans*. The orange arrows represent DMA and methanethiol (MeSH) specific reactions that catalyzes the transfer of the methyl groups from DMS to ultimately form methyl-CoM (CH3-CoM). CH3-CoM undergoes a disproportionation reaction to generate CO2 and CH4 in a 3:1 ratio.

**Supplementary Figure 9:**
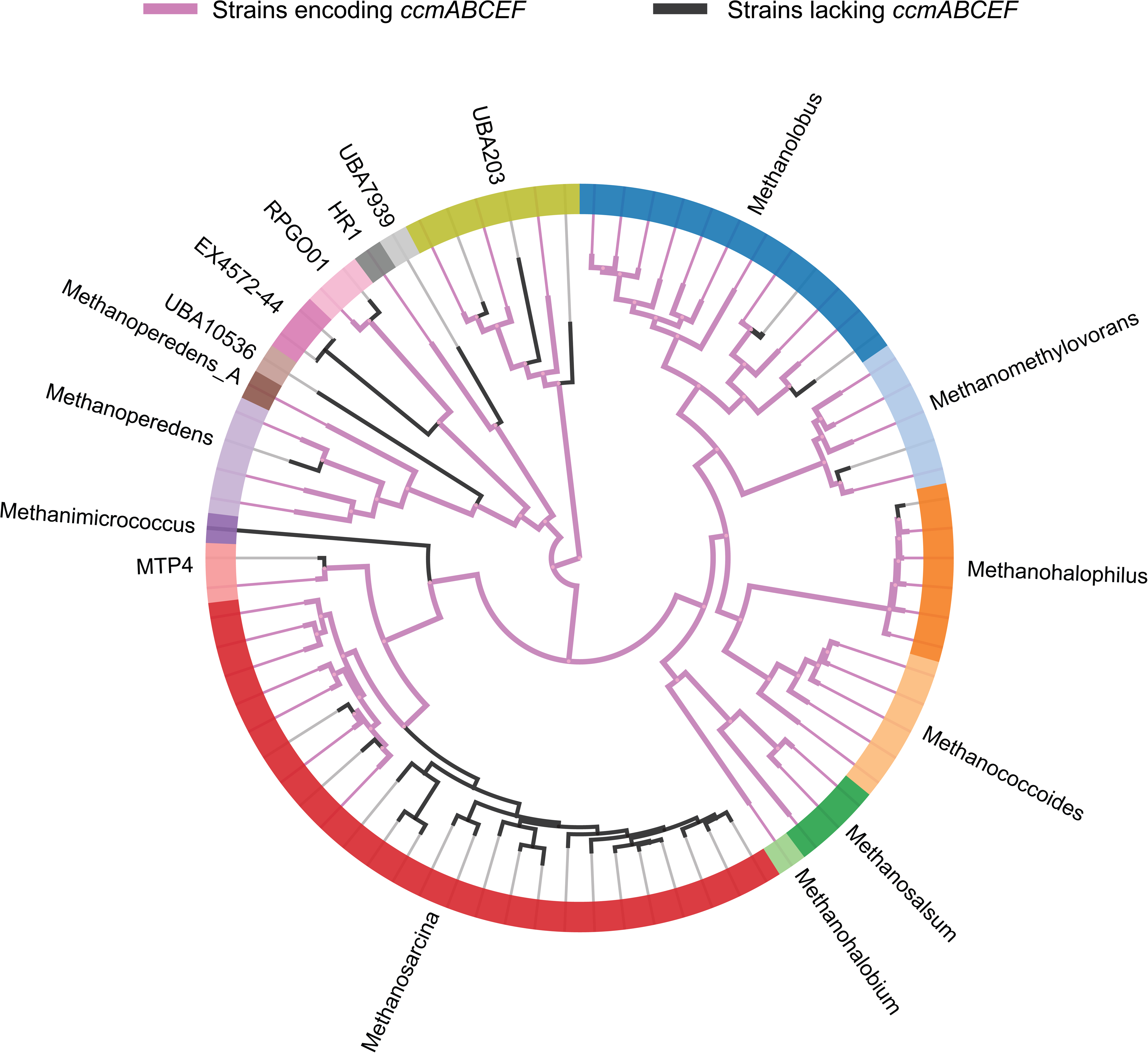
A phylogenomic reconstruction of the relationship between various genera (depicted by different colors along the circumference of the radial tree) within the Family *Methanosarcinaceae* from Annotree (Mendler *et al.*, 2019). Purple lines depict strains that encode homologs of *ccmABCEF1F2* in their genome whereas black lines depict strains that lack homologs of *ccmABCEF1F2* in their genome. Based on the pattern of distribution of the *ccmABCEF1F2* genes it is likely that these genes were present in the last common ancestor of the *Methanosarcinaceae* and have been lost in members of various Genera including *Methanosarcina* spp.*, Methanohalophilus* spp*., Methanomethylovorans* spp.*, Methanolobus* spp. and *Methanoperedens* spp.

**Supplementary Figure 10:**
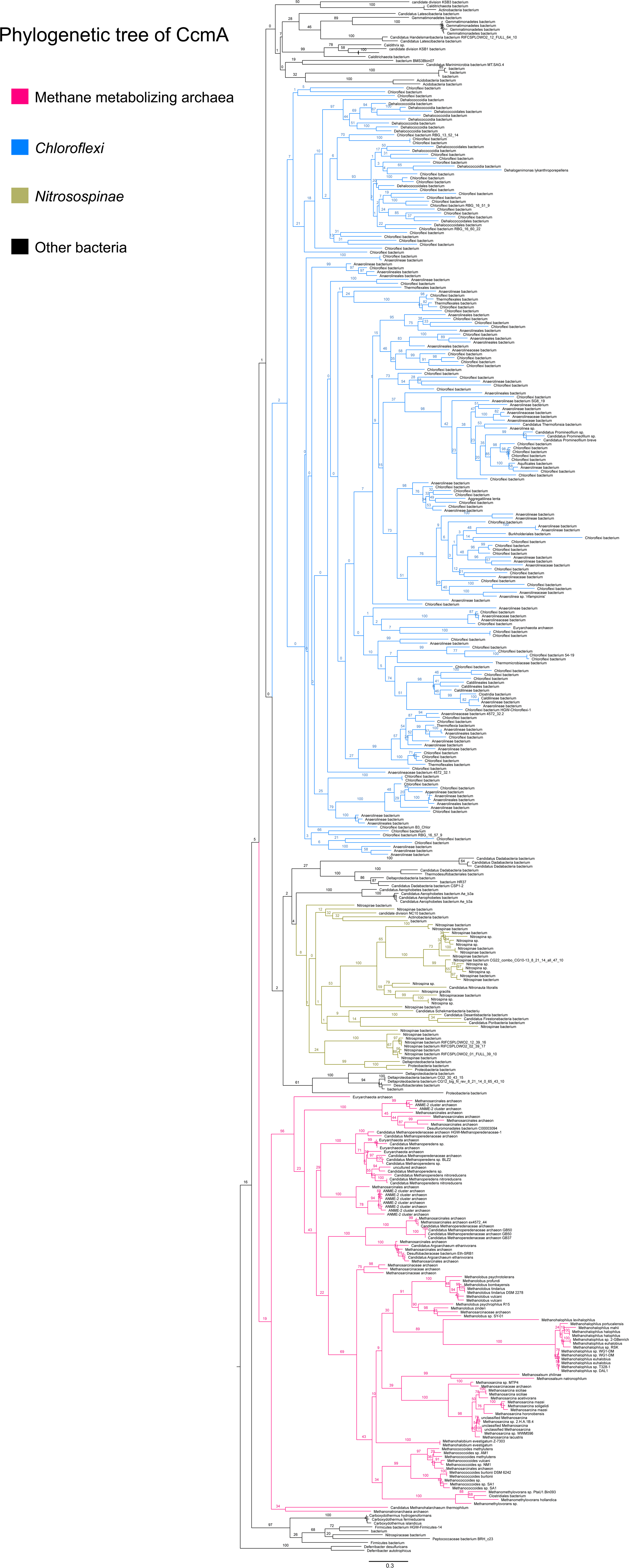
An unrooted maximum-likelihood tree of 500 CcmA sequences retrieved from the NCBI non-redundant protein sequence database using the corresponding sequence from *Methanosarcina acetivorans* as the search query. Branch labels indicate support values.

**Supplementary Figure 11:**
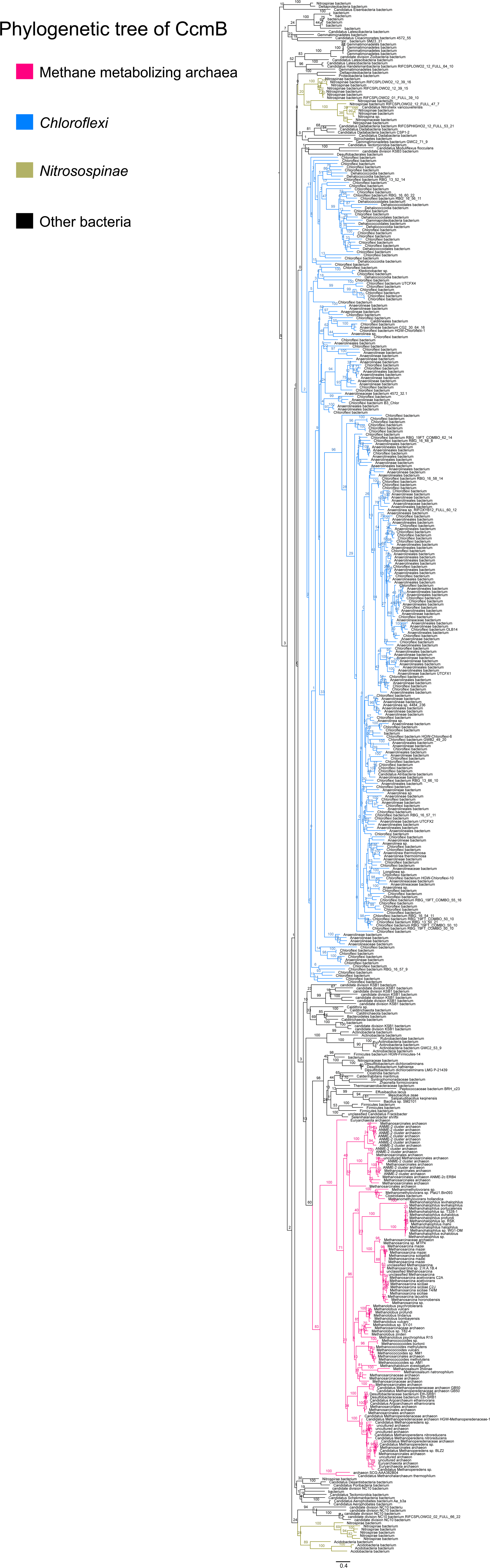
An unrooted maximum-likelihood tree of 500 CcmB sequences retrieved from the NCBI non-redundant protein sequence database using the corresponding sequence from *Methanosarcina acetivorans* as the search query. Branch labels indicate support values.

**Supplementary Figure 12:**
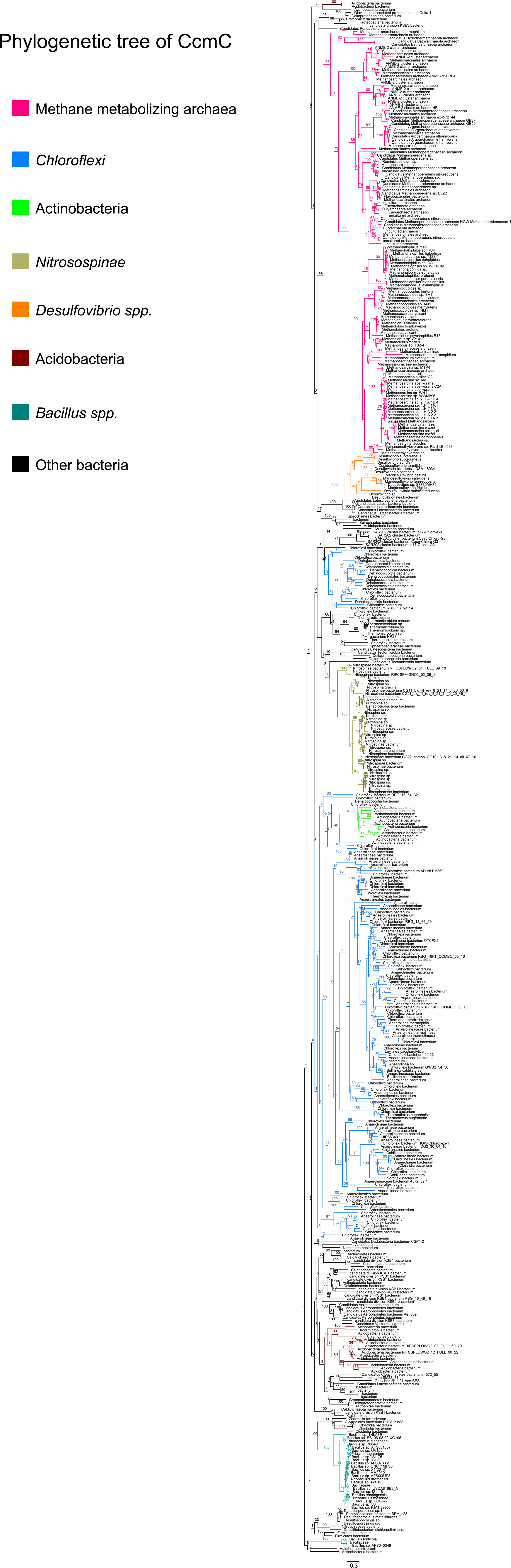
An unrooted maximum-likelihood tree of 500 CcmC sequences retrieved from the NCBI non-redundant protein sequence database using the corresponding sequence from *Methanosarcina acetivorans* as the search query. Branch labels indicate support values.

**Supplementary Figure 13:**
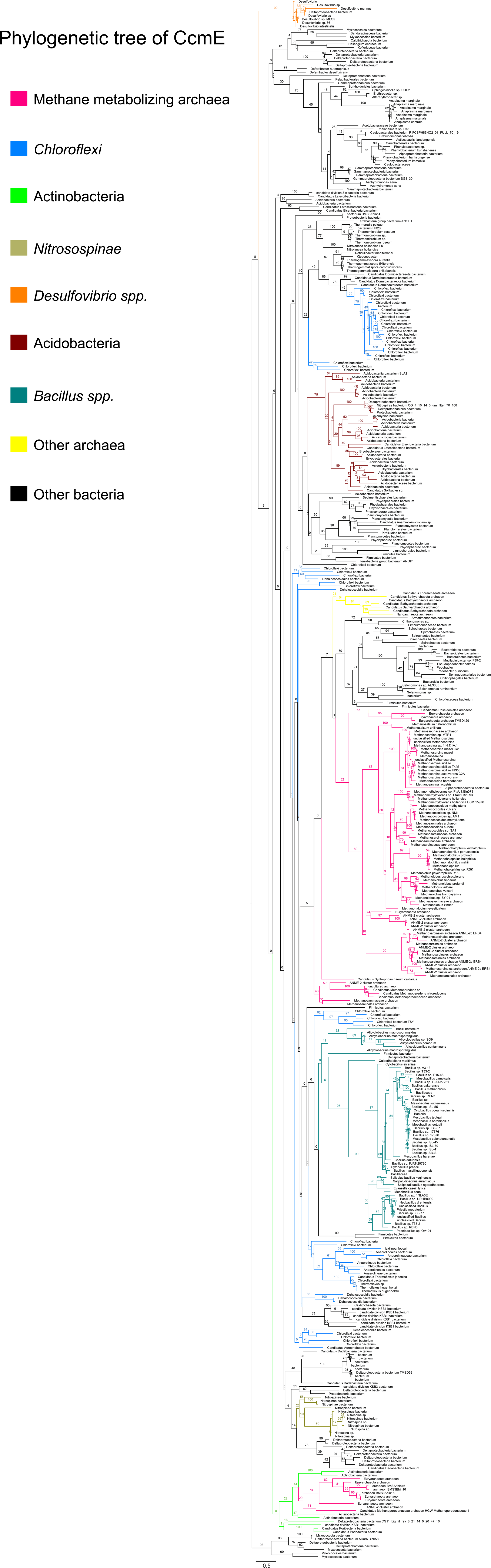
An unrooted maximum-likelihood tree of 500 CcmE sequences retrieved from the NCBI non-redundant protein sequence database using the corresponding sequence from *Methanosarcina acetivorans* as the search query. Branch labels indicate support values.

**Supplementary Figure 14:**
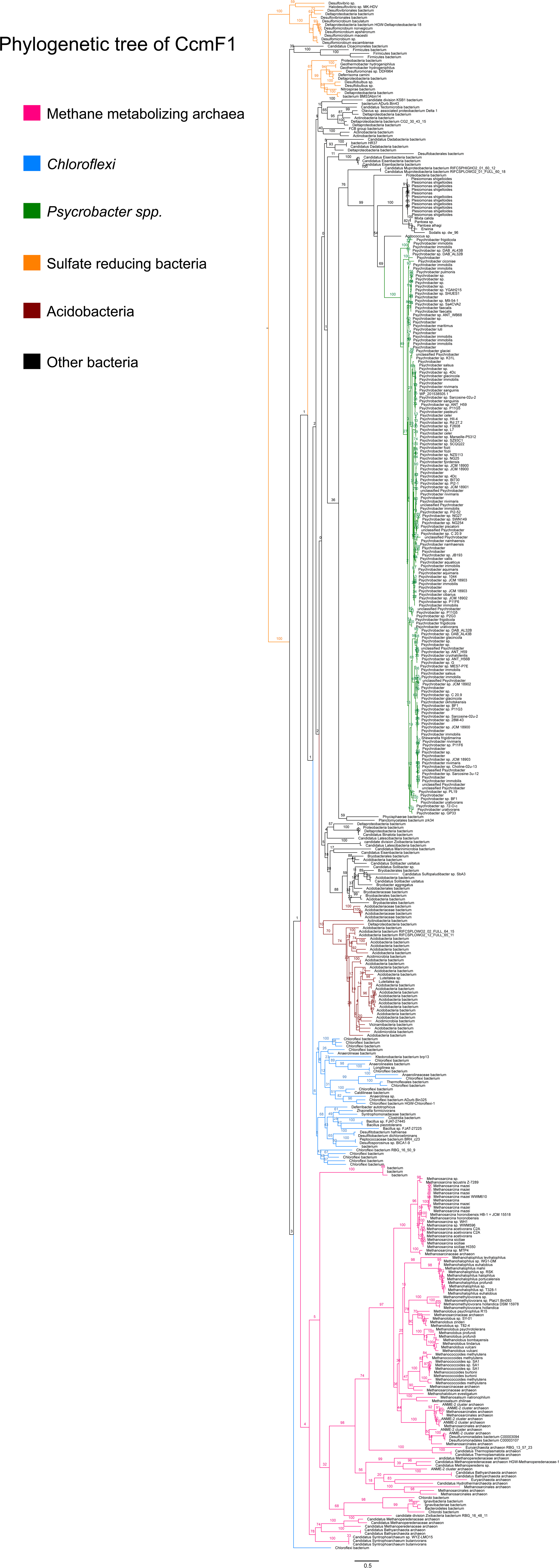
An unrooted maximum-likelihood tree of 500 CcmF1 sequences retrieved from the NCBI nonredundant protein sequence database using the corresponding sequence from *Methanosarcina acetivorans* as the search query. Branch labels indicate support values.

**Supplementary Figure 15:**
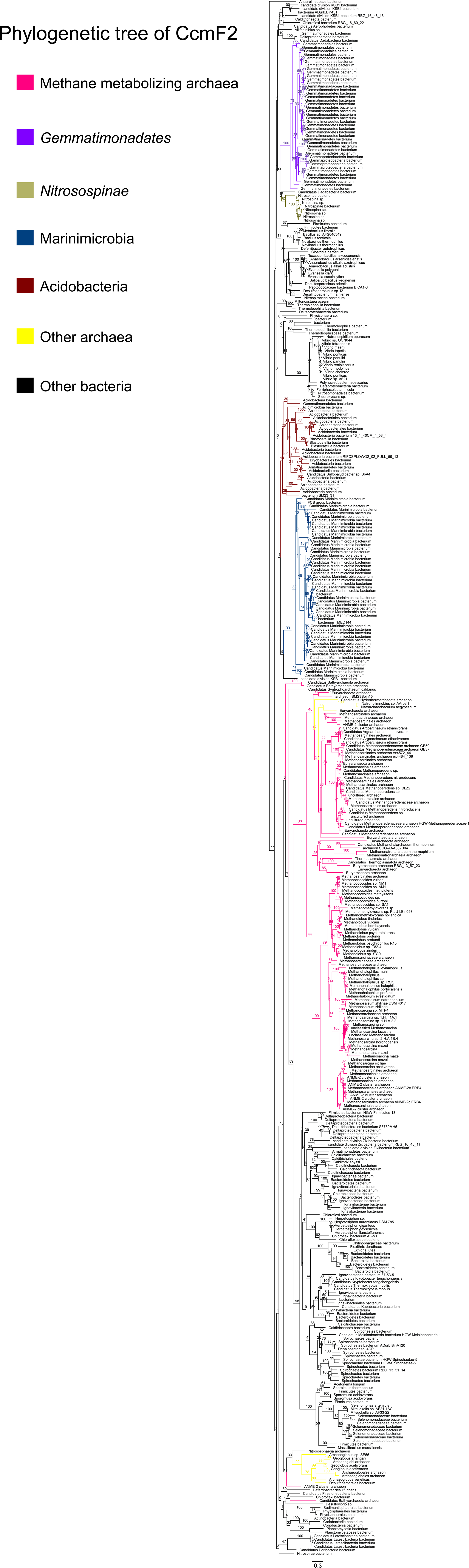
An unrooted maximum-likelihood tree of 500 CcmF2 sequences retrieved from the NCBI non-redundant protein sequence database using the corresponding sequence from *Methanosarcina acetivorans* as the search query. Branch labels indicate support values.

**Supplementary Table 1:** List of mutations in CRISPR-edited mutant strain DDN029 containing a Δ *ccmABC* in-frame deletion mutation

| Position | Mutation | Notes |
| --- | --- | --- |
| 487691 | C added | Present in Parent (WWM60) |
| 836182-836771 | $\Delta hpt::PmcrB-tetR$ | Present in Parent (WWM60) |
| 941168 | A5->6 | Present in Parent (WWM60) |
| 1314120 | $\Delta 1bp$ | Present in Parent (WWM60) |
| **1698001** | **$\Delta 2684 bp$** | **$\Delta ccmABC$ (only present in DDN029)** |
| 2086881-82 | TC->CT | Present in Parent (WWM60) |
| 2086886 | G->T | Present in Parent (WWM60) |
| 2534543 | C added | Present in Parent (WWM60) |
| 2836646 | A->G (F64L) | Present in Parent (WWM60) |
| 2867059 | T->G (H313Q) | Present in Parent (WWM60) |
| 3433201 | $\Delta 1bp$ | Present in Parent (WWM60) |
| 4295452 | $\Delta 1bp$ | Present in Parent (WWM60) |
| 4874567 | $\Delta 1bp$ | Present in Parent (WWM60) |
| 4945345 | C added | Present in Parent (WWM60) |
| 5078585 | G->A (M1M) | Present in Parent (WWM60) |

**Supplementary Table 2:**
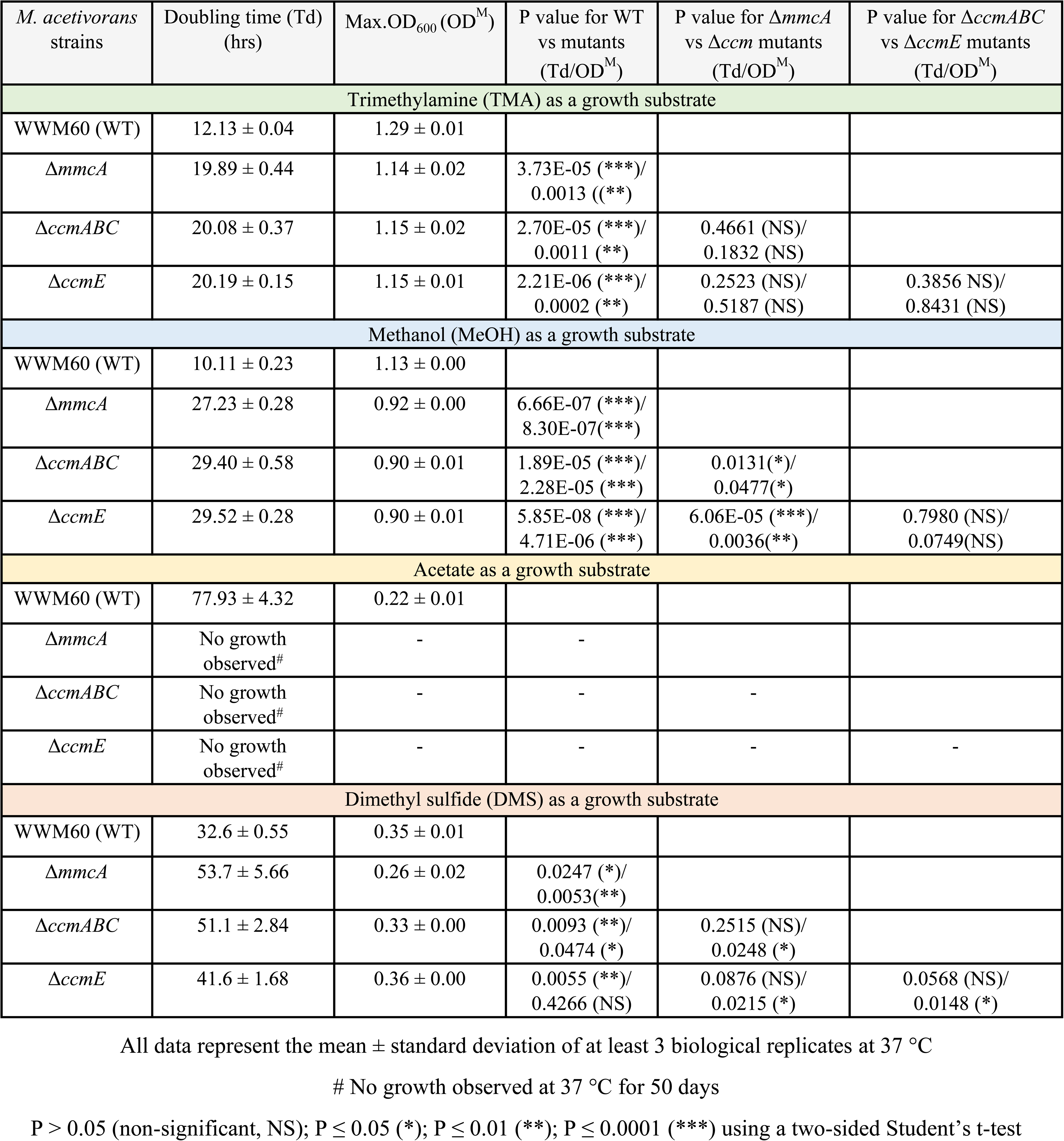
Growth data of *M. acetivorans* strains shown in Figure 6

**Supplementary Table 3:**
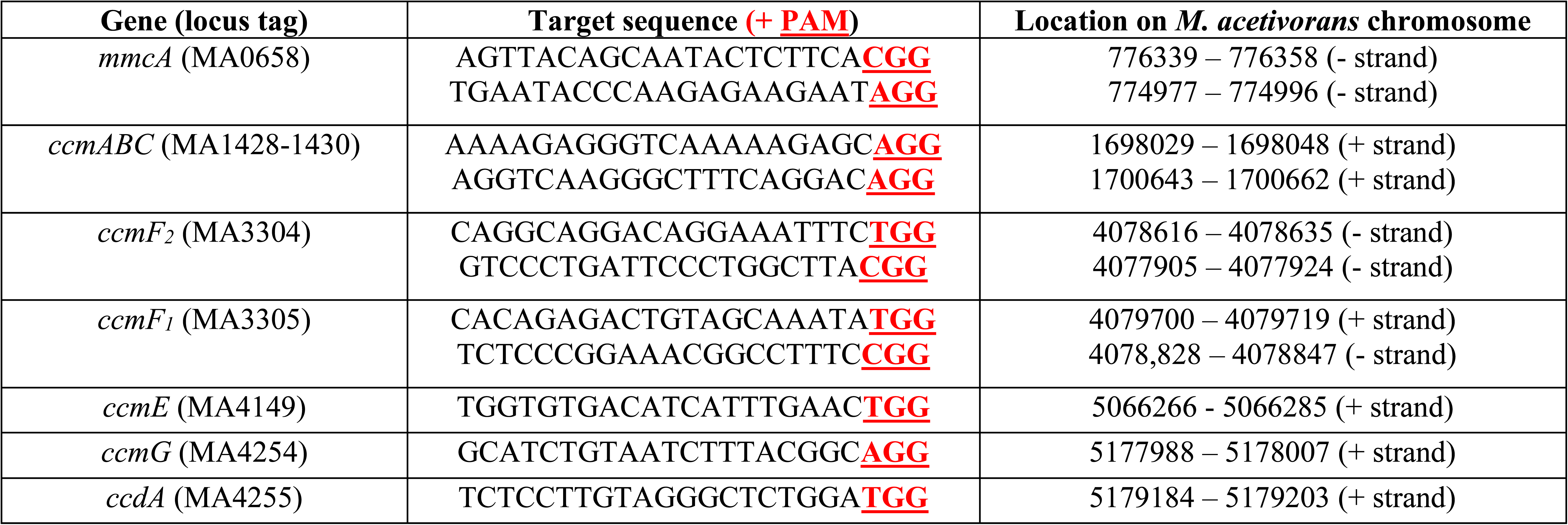
List of target sequences used in this study

**Supplementary Table 4:** List of plasmids used in this study

| Plasmid | Features | Source |
| --- | --- | --- |
| pAMG40 | Vector for fosmid retrofitting that contains pC2A and λattB | (Guss et al. 2008) |
| pJK027A | Vector with *Pmc*r*B*(*tet*O1) promoter fusion to *uidA* that contains ϕC31-attB and λattP | (Guss et al. 2008) |
| pJK029A | Vector with *Pmc*r*B*(*tet*O4) promoter fusion to *uidA* that contains ϕC31-attB and λattP | (Guss et al. 2008) |
| pDN201 | pJK027A-derived plasmid with *Pmc*r*B*(*tet*O1) promoter fusion to *Spy cas*9 | (Nayak and Metcalf, 2017) |
| pDN401 | pDN201-derived plasmid with synthetic fragments containing *PmtaCB1* promoter fusion to sgRNAs in tandem targeting the *mmcA* (MA0658) coding sequence (CDS) | This study |
| pDN402 | pDN401-derived plasmid containing a repair template to generate an in-frame deletion of the *mmcA* (MA0658) CDS | This study |
| pDN403 | Cointegrate of pDN401 and pAMG40 | This study |
| pDN404 | Cointegrate of pDN402 and pAMG40 | This study |
| pDN406 | pJK029A-derived plasmid with *Pmc*r*B*(*tet*O4) promoter fusion to *mmcA* (MA0658) CDS from *M. acetivorans* with a C-terminal tandem affinity purification tag (containing a 3X FLAG tag and a Twin-Strep tag) | This study |
| pDN409 | Cointegrate of pDN406 and pAMG40 | This study |
| pDN411 | pDN201-derived plasmid with synthetic fragments containing *PmtaCB1* promoter fusion to sgRNAs in tandem targeting the *ccmE* (MA4149) coding sequence (CDS) | This study |
| pDN412 | pDN411-derived plasmid containing a repair template to generate an in-frame deletion of the *ccmE* (MA4149) CDS | This study |
| pDN413 | Cointegrate of pDN411 and pAMG40 | This study |
| pDN414 | Cointegrate of pDN412 and pAMG40 | This study |
| pDN421 | pDN201-derived plasmid with synthetic fragments containing *PmtaCB1* promoter fusion to sgRNAs in tandem targeting the *ccmF1* (MA3305) coding sequence (CDS) | This study |
| pDN422 | pDN201-derived plasmid with synthetic fragments containing *PmtaCB1* promoter fusion to sgRNAs in tandem targeting the *ccmF2* (MA3304) coding sequence (CDS) | This study |
| pDN423 | pDN201-derived plasmid with synthetic fragments containing *PmtaCB1* promoter fusion to sgRNAs in tandem targeting the *ccmF1F2* (MA3305-3304) coding sequences (CDS) | This study |
| pDN424 | pDN421-derived plasmid containing a repair template to generate an in-frame deletion of the *ccmF1* (MA3305) CDS | This study |
| pDN425 | pDN422-derived plasmid containing a repair template to generate an in-frame deletion of the *ccmF2* (MA3304) CDS | This study |
| pDN426 | pDN423-derived plasmid containing a repair template to generate an in-frame deletion of the *ccmF1F2* (MA3305-3304) CDS | This study |
| pDN427 | Cointegrate of pDN421 and pAMG40 | This study |
| pDN428 | Cointegrate of pDN422 and pAMG40 | This study |
| pDN429 | Cointegrate of pDN423 and pAMG40 | This study |
| pDN430 | Cointegrate of pDN424 and pAMG40 | This study |
| pDN431 | Cointegrate of pDN425 and pAMG40 | This study |
| pDN432 | Cointegrate of pDN426 and pAMG40 | This study |
| pDN443 | pDN201-derived plasmid with synthetic fragments containing *PmtaCB1* promoter fusion to sgRNAs in tandem targeting the *ccmABC* (MA1428-1430) coding sequences (CDS) | This study |
| pDN444 | pDN443-derived plasmid containing a repair template to generate an in-frame deletion of the *ccmABC* (MA1428-1430) CDS | This study |
| pDN445 | Cointegrate of pDN443 and pAMG40 | This study |
| pDN446 | Cointegrate of pDN444 and pAMG40 | This study |
| pDPG001 | pDN201-derived plasmid with synthetic fragments containing *PmtaCB1* promoter fusion to sgRNAs in tandem targeting the *ccmG* (MA4254) coding sequence (CDS) | This study |
| pDPG002 | pDN201-derived plasmid with synthetic fragments containing *PmtaCB1* promoter fusion to sgRNAs in tandem targeting the *ccdA* (MA4255) coding sequence (CDS) | This study |
| pDPG003 | pDN201-derived plasmid with synthetic fragments containing *PmtaCB1* promoter fusion to sgRNAs in tandem targeting the *ccmG-ccdA* (MA4254-4255) coding sequences (CDS) | This study |
| pDPG004 | pDPG001-derived plasmid containing a repair template to generate an in-frame deletion of the *ccmG* (MA4254) CDS | This study |
| pDPG005 | pDPG002-derived plasmid containing a repair template to generate an in-frame deletion of the *ccdA* (MA4255) CDS | This study |
| pDPG006 | pDPG003-derived plasmid containing a repair template to generate an in-frame deletion of the *ccmG-ccdA* (MA4254-4254) CDS | This study |
| pDPG007 | Cointegrate of pDPG004 and pAMG40 | This study |
| pDPG008 | Cointegrate of pDPG005 and pAMG40 | This study |
| pDPG009 | Cointegrate of pDPG006 and pAMG40 | This study |
| pDPG010 | Cointegrate of pJK029A and pAMG40 | This study |
| pDPG014 | pJK029A-derived plasmid with *PmcrB(tetO4)* promoter fusion to *ccmABC* (MA1428-1430)-*ccmE* (MA4149)-*ccmF1F2*(MA3305-3304) CDS from *M. acetivorans* | This study |
| pDPG015 | pJK029A-derived plasmid with *M. barkeri* *PserC*(constitutive) promoter fusion to *mmcA* (MA0658) CDS from *M. acetivorans* with a C-terminal tandem affinity purification tag (containing a 3X FLAG tag and a Twin-Strep sequence) | This study |
| pDPG017 | pJK029A-derived plasmid with *PmcrB(tetO4)* promoter fusion to *ccmABC* (MA1428-1430)-*ccmE* (MA4149)-*ccmF1F2*(MA3305-3304) CDS from *M. acetivorans* and *M. barkeri* *PserC*(constitutive) promoter fusion to *mmcA* (MA0658) CDS from *M. acetivorans* with a C-terminal tandem affinity purification tag (containing a 3X FLAG tag and a Twin-Strep sequence) | This study |
| pKES30 | pJK029A-derived plasmid with *PmcrB(tetO4)* promoter fusion to *ccmE* (MA4149) CDS from *M. acetivorans* with a C-terminal tandem affinity purification tag (containing a 1X Strep and a 1X FLAG sequence) | This study |
| pKES031 | Cointegrate of pKES30 and pAMG40 | This study |
| pKES038 | pJK029A-derived plasmid with *PmcrB(tetO4)* promoter fusion to *ccmE* (MA4149) CDS with C120H substitution with a C-terminal tandem affinity purification tag (containing a 1X Strep and a 1X FLAG sequence) | This study |
| pKES039 | Cointegrate of pKES38 and pAMG40 | This study |
| pKES040 | pJK029A-derived plasmid with *PmcrB(tetO4)* promoter fusion to *ccmE* (MA4149) CDS with C120A substitution with a C-terminal tandem affinity purification tag (containing a 1X Strep and a 1X FLAG sequence) | This study |
| pKES041 | Cointegrate of pKES40 and pAMG40 | This study |

Note: pDN201/pJK029A derived plasmids contain the chloramphenicol resistance marker, pAMG40 contains kanamycin resistance marker, and the co-integrates of pDN201/pJK029A derived plasmids and pAMG40 contain both contain chloramphenicol and kanamycin resistance markers.

**Supplementary Table 5:**
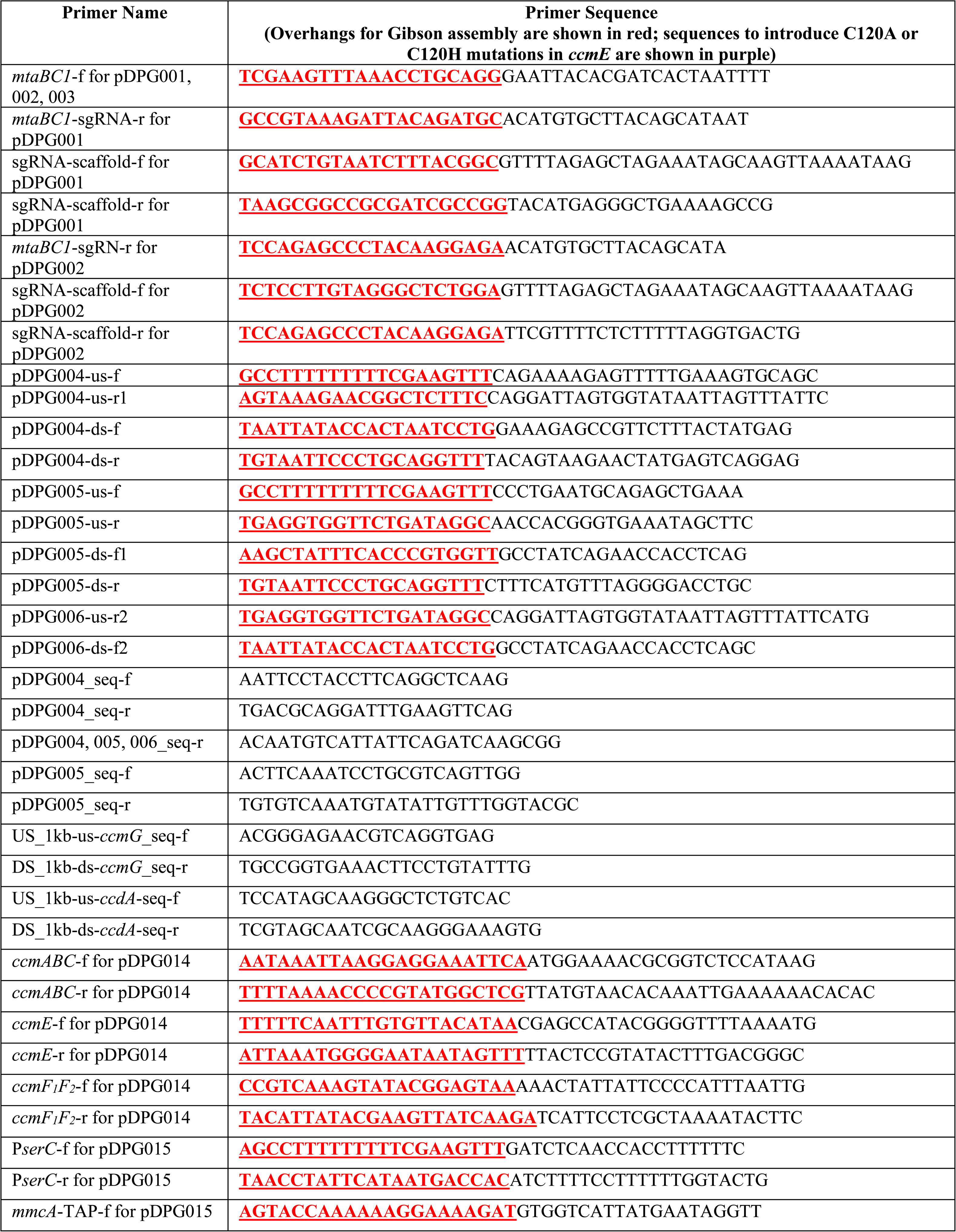
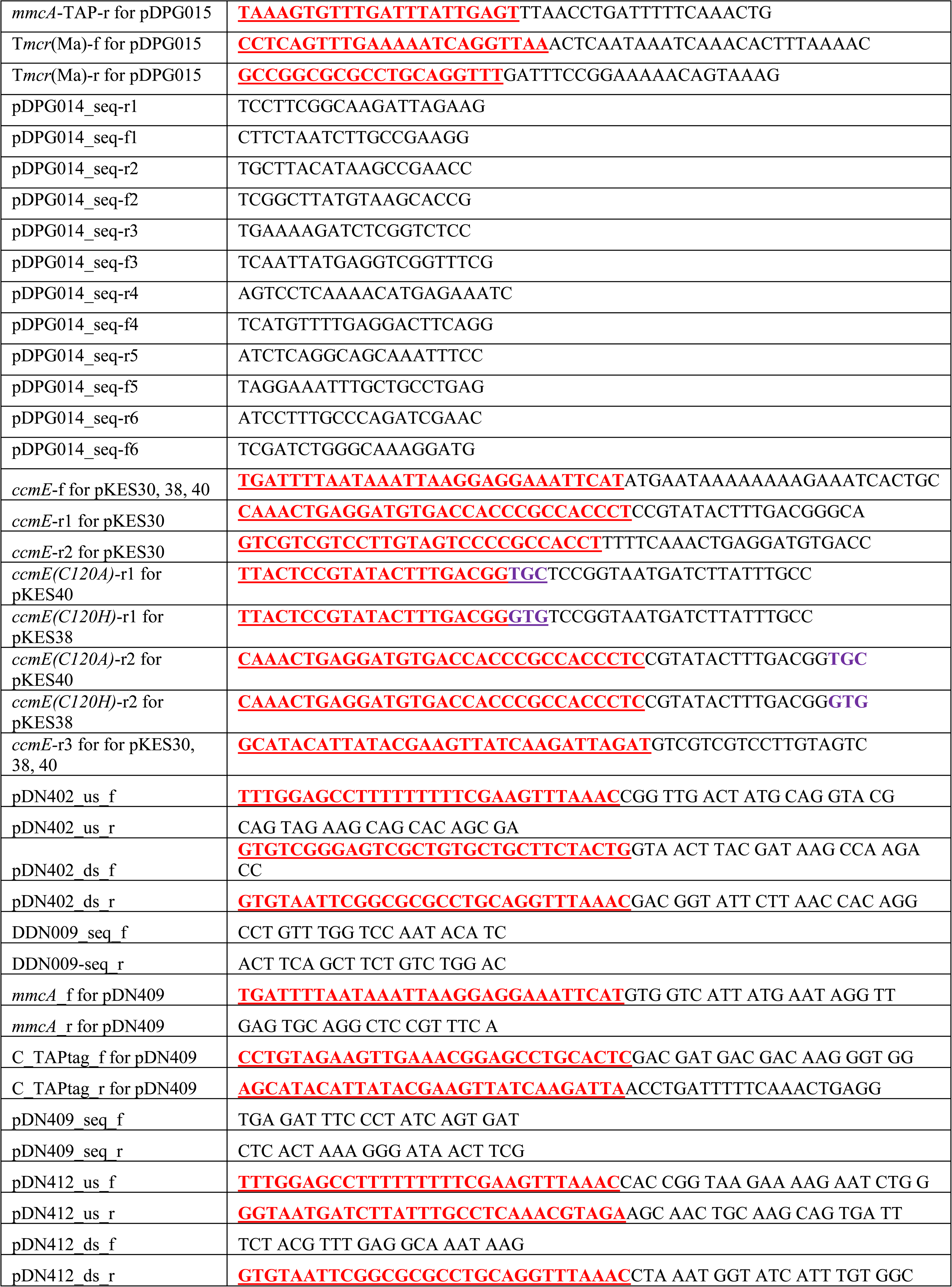
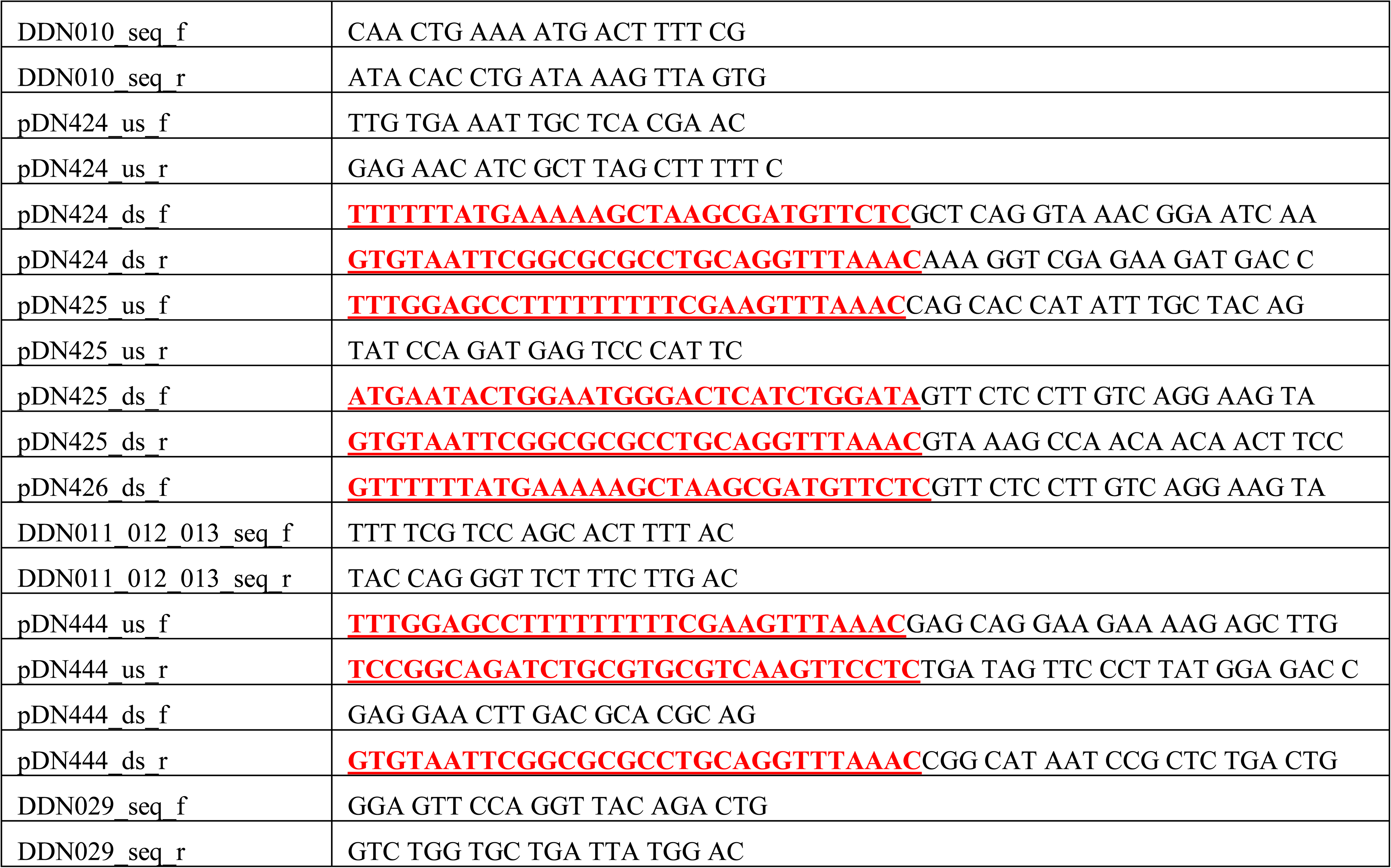
List of primers used in this study

**Supplementary Table 6:**
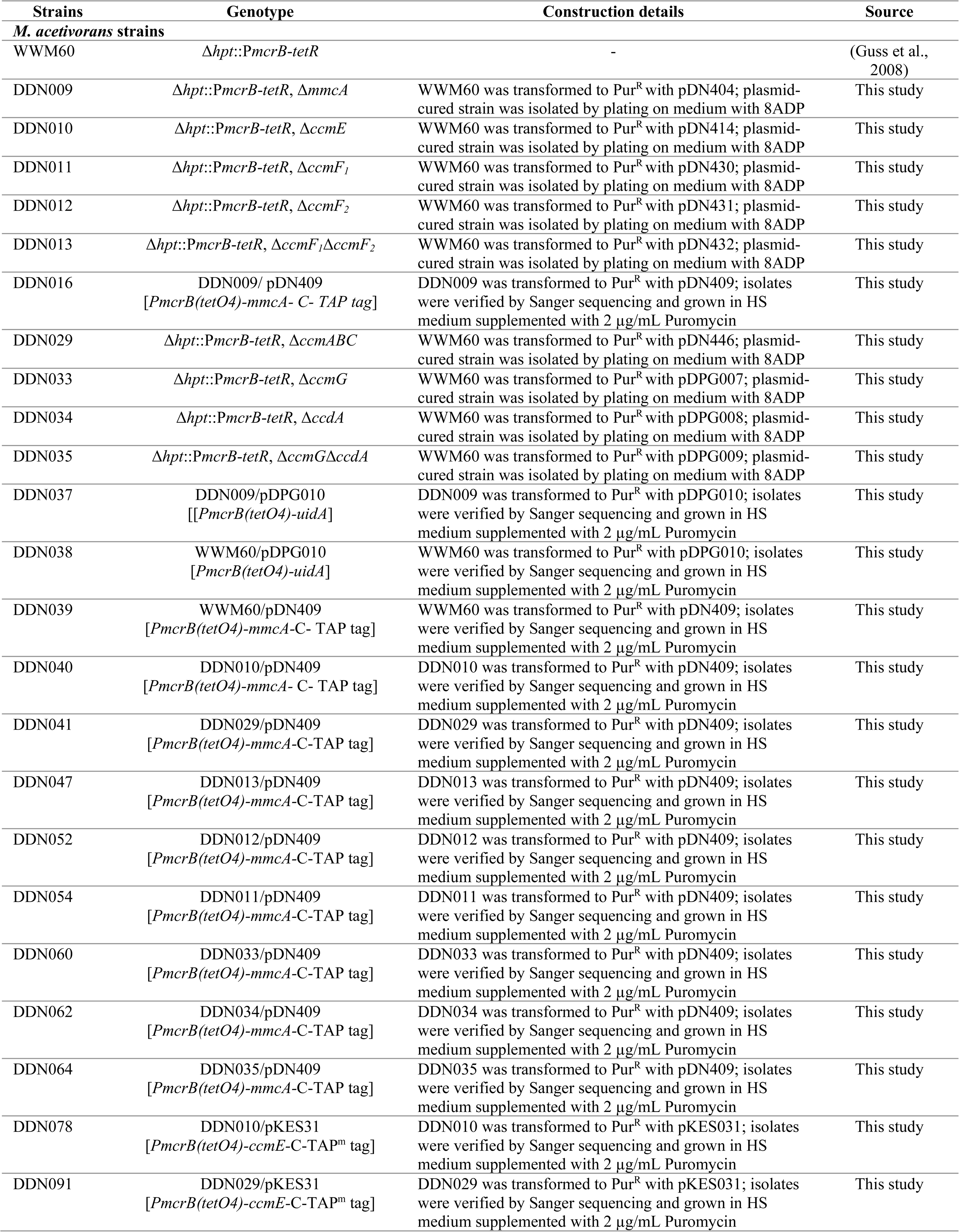
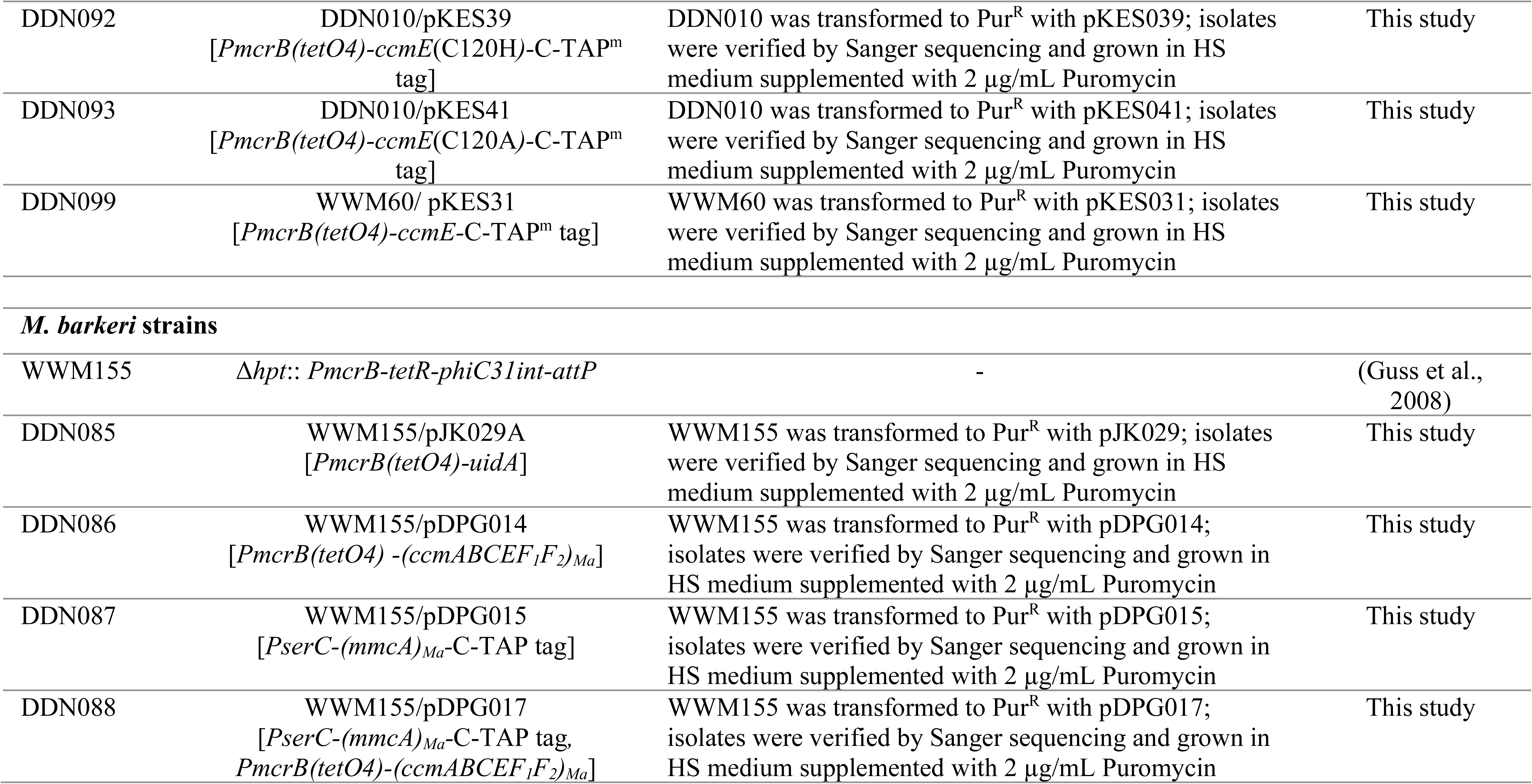
List of *Methanosarcina* strains used in this study

## References

1. Richardson DJ. Bacterial respiration: a flexible process for a changing environment. Microbiology 2000; 146:551–557

2. Bertini I, Cavallaro G, Rosato A. Cytochrome c: Occurrence and functions. Chemical Reviews 2006: 106:90–115

3. Shi L, Dong H, Reguera G, Beyenal H, Lu A, Liu J, et al. Extracellular electron transfer mechanisms between microorganisms and minerals. Nature Reviews Microbiology 2016; 14:651–662

4. Gupta D, Guzman MS, Bose A. Extracellular electron uptake by autotrophic microbes: physiological, ecological, and evolutionary implications. Journal of Industrial Microbiology and Biotechnology 2020; 47:863–876

5. Kranz RG, Richard-Fogal C, Taylor J-S, Frawley ER. Cytochrome c Biogenesis: Mechanisms for Covalent Modifications and Trafficking of Heme and for Heme-Iron Redox Control. Microbiology and Molecular Biology Reviews 2009; 73:510–528

6. Sanders C, Turkarslan S, Lee DW, Daldal F. Cytochrome c biogenesis: The Ccm system. Trends in Microbiology 2010; 18:266–274

7. Sambongi Y, Stoll R, Ferguson SJ. Alteration of haem-attachment and signal-cleavage sites for *Paracoccus denitrificans* cytochrome C550 probes pathway of c-type cytochrome biogenesis in *Escherichia coli*. Molecular Microbiology 1996; 19:1193–1204

8. Thöny-Meyer L, Künzler P. Translocation to the periplasm and signal sequence cleavage of preapocytochrome c depend on sec and lep, but not on the ccm gene products. European Journal of Biochemistry 1997; 246:794–799

9. Helde R, Wieseler B, Wachter E, Neubüser A, Hoffschulte HK, Hengelage T, et al. Comparative characterization of secA from the α-subclass purple bacterium *Rhodobacter capsulatus* and *Escherichia coli* reveals differences in membrane and precursor specificity. Journal of Bacteriology 1997; 179:4003–4012

10. Kranz R, Lill R, Goldman B, Bonnard G, Merchant S. Molecular mechanisms of cytochrome c biogenesis: Three distinct systems. Molecular Microbiology 1998; 29:383–396

11. Allen JWA, Harvat EM, Stevens JM, Ferguson SJ. A variant System I for cytochrome c biogenesis in archaea and some bacteria has a novel CcmE and no CcmH. FEBS Letters 2006; 580:4827–4834

12. Kletzin A, Heimerl T, Flechsler J, Niftrik L van, Rachel R, Klingl A. Cytochromes c in Archaea: Distribution, maturation, cell architecture, and the special case of *Ignicoccus hospitalis*. Frontiers in Microbiology 2015; 6:439

13. Chadwick GL, Skennerton CT, Laso-Perez R, Leu AO, Speth DR, Yu H, et al. Comparative genomics reveals electron transfer and syntrophic mechanisms differentiating methanotrophic and methanogenic archaea. bioRxiv 2021.

14. Giegé P, Grienenberger JM, Bonnard G. Cytochrome c biogenesis in mitochondria. Mitochondrion 2008; 8:61–73

15. Fabianek RA, Hennecke H, Thöny-Meyer L. Periplasmic protein thiol:disulfide oxidoreductases of *Escherichia coli*. FEMS Microbiology Reviews 2000; 24:303–316

16. Setterdahl AT, Goldman BS, Hirasawa M, Jacquot P, Smith AJ, Kranz RG, et al. Oxidation - Reduction properties of disulfide-containing proteins of the *Rhodobacter capsulatus* cytochrome c biogenesis system. Biochemistry 2000; 39:10172–10176

17. Reid E, Cole J, Eaves DJ. The *Escherichia coli* CcmG protein fulfils a specific role in cytochrome c assembly. Biochemical Journal 2001; 355:51–58

18. Thöny-Meyer L. Biogenesis of respiratory cytochromes in bacteria. Microbiology and Molecular Biology Reviews 1997; 61:337–376

19. Goddard AD, Stevens JM, Rao F, Mavridou DAI, Chan W, Richardson DJ, et al. c-Type cytochrome biogenesis can occur via a natural Ccm system lacking CcmH, CcmG, and the heme-binding histidine of CcmE. Journal of Biological Chemistry 2010; 285:22882–22889

20. Fu H, Jin M, Wan F, Gao H. *Shewanella oneidensis* cytochrome c maturation component CcmI is essential for heme attachment at the non-canonical motif of nitrite reductase NrfA. Molecular Microbiology 2015; 95:410–425

21. Richard-Fogal CL, Frawley ER, Kranz RG. Topology and function of CcmD in cytochrome C maturation. Journal of Bacteriology 2008; 190:3489–3493

22. Sreeramulu K. Purification and partial characterization of cytochrome c 552 from Halobacterium salinarium. Indian Journal of Biochemistry and Biophysics 2003; 40:274–277

23. Naß B, Pöll U, Langer JD, Kreuter L, Küper U, Flechsler J, et al. Three multihaem cytochromes c from the hyperthermophilic archaeon *Ignicoccus hospitalis*: Purification, properties and localization. Microbiology 2014; 160:1278–1289

24. Smith JA, Aklujkar M, Risso C, Leang C, Giloteaux L, Holmes DE. Mechanisms involved in Fe(III) respiration by the hyperthermophilic archaeon *Ferroglobus placidus*. Applied and Environmental Microbiology 2015; 81:2735–2744

25. Feinberg LF, Srikanth R, Vachet RW, Holden JF. Constraints on anaerobic respiration in the hyperthermophilic archaea *Pyrobaculum islandicum* and *Pyrobaculum aerophilum*. Applied and Environmental Microbiology 2008; 74:396–402

26. Wang M, Tomb JF, Ferry JG. Electron transport in acetate-grown *Methanosarcina acetivorans*. BMC Microbiology 2011; 11:165

27. Arshad A, Speth DR, de Graaf RM, Op den Camp HJM, Jetten MSM, Welte CU. A metagenomics-based metabolic model of nitrate-dependent anaerobic oxidation of methane by *Methanoperedens*-like archaea. Frontiers in Microbiology 2015; 6:1423

28. Holmes DE, Ueki T, Tang HY, Zhou J, Smith JA, Chaput G, et al. A membrane-bound cytochrome enables *Methanosarcina acetivorans* to conserve energy from extracellular electron transfer. mBio 2019; 10:e00789–19

29. Sowers KR, Boone JE, Gunsalus RP. Disaggregation of *Methanosarcina spp.* and growth as single cells at elevated osmolarity. Applied and Environmental Microbiology 1993; 59:3832–3839

30. Guss AM, Rother M, Zhang JK, Kulkarni G, Metcalf WW. New methods for tightly regulated gene expression and highly efficient chromosomal integration of cloned genes for *Methanosarcina* species. Archaea 2008; 2:193–203

31. Kohler PRA, Metcalf WW. Genetic manipulation of *Methanosarcina spp*. Frontiers in Microbiology 2012; 3:259

32. Nayak DD, Metcalf WW. Cas9-mediated genome editing in the methanogenic archaeon Methanosarcina acetivorans. Proceedings of the National Academy of Sciences of the United States of America 2017; 114:2976–2981

33. Li Q, Li L, Rejtar T, Lessner DJ, Karger BL, Ferry JG. Electron transport in the pathway of acetate conversion to methane in the marine archaeon *Methanosarcina acetivorans*. Journal of Bacteriology 2006; 188:702–710

34. Schlegel K, Welte C, Deppenmeier U, Müller V. Electron transport during aceticlastic methanogenesis by *Methanosarcina acetivorans* involves a sodium-translocating Rnf complex. FEBS Journal 2012; 279:4444–4452

35. Ferry JG. *Methanosarcina acetivorans*: A Model for Mechanistic Understanding of Aceticlastic and Reverse Methanogenesis. Frontiers in Microbiology 2020; 11:1806

36. Holmes DE, Zhou J, Ueki T, Woodard T, Lovley DR. Mechanisms for Electron Uptake by *Methanosarcina acetivorans* during Direct Interspecies Electron Transfer. mBio 2021; 12:e0234421

37. Nayak DD, Mahanta N, Mitchell DA, Metcalf WW. Post-translational thioamidation of methyl-coenzyme M reductase, a key enzyme in methanogenic and methanotrophic archaea. eLife 2017; 6:e29218

38. Nayak DD, Liu A, Agrawal N, Rodriguez-Carerro R, Dong SH, Mitchell DA, et al. Functional interactions between posttranslationally modified amino acids of methyl-coenzyme M reductase in *Methanosarcina acetivorans*. PLoS Biology 2020; 18:e3000507

39. Zhou J, Holmes DE, Tang HY, Lovley DR. Correlation of Key Physiological Properties of *Methanosarcina* Isolates with Environment of Origin. Applied and Environmental Microbiology 2021; 87:e00731–21

40. Borrel G, Adam PS, McKay LJ, Chen LX, Sierra-García IN, Sieber CMK, et al. Wide diversity of methane and short-chain alkane metabolisms in uncultured archaea. Nature Microbiology 2019; 4:603–613

41. Feissner R, Xiang Y, Kranz RG. Chemiluminescent-based methods to detect subpicomole levels of c-type cytochromes. Analytical Biochemistry 2003; 315:90–94

42. Goldman BS, Beckman DL, Bali A, Monika EM, Gabbert KK, Kranz RG. Molecular and immunological analysis of an ABC transporter complex required for cytochrome c biogenesis. Journal of Molecular Biology 1997; 268:724–738

43. Feissner RE, Richard-Fogal CL, Frawley ER, Loughman JA, Earley KW, Kranz RG. Recombinant cytochromes c biogenesis systems I and II and analysis of haem delivery pathways in *Escherichia coli*. Molecular Microbiology 2006; 60:563–577

44. Allen JWA, Ginger ML, Ferguson SJ. Complexity and diversity in c-type cytochrome biognenesis systems. Biochemical Society Transactions 2005;33:145–146

45. Christensen O, Harvat EM, Thöny-Meyer L, Ferguson SJ, Stevens JM. Loss of ATP hydrolysis activity by CcmAB results in loss of c-type cytochrome synthesis and incomplete processing of CcmE. FEBS Journal 2007; 274:2322–2332

46. Feissner RE, Richard-Fogal CL, Frawley ER, Kranz RG. ABC transporter-mediated release of a haem chaperone allows cytochrome c biogenesis. Molecular Microbiology 2006; 61:219–231

47. Schulz H, Hennecke H, Thöny-Meyer L. Prototype of a heme chaperone essential for cytochrome c maturation. Science 1998; 281:1197–1200

48. Parks DH, Chuvochina M, Rinke C, Mussig AJ, Chaumeil P-A, Hugenholtz P. GTDB: an ongoing census of bacterial and archaeal diversity through a phylogenetically consistent, rank normalized and complete genome-based taxonomy. Nucleic Acids Research 2021;gkab776

49. Mendler K, Chen H, Parks DH, Lobb B, Hug LA, Doxey AC. Annotree: Visualization and exploration of a functionally annotated microbial tree of life. Nucleic Acids Research 2019; 47: 4442–4448

50. Enggist E, Thöny-Meyer L. The C-terminal flexible domain of the heme chaperone CcmE is important but not essential for its function. Journal of Bacteriology 2003; 185:3821–3827

51. Stevens JM, Daltrop O, Higham CW, Ferguson SJ. Interaction of heme with variants of the heme chaperone CcmE carrying active site mutations and a cleavable N-terminal his tag. Journal of Biological Chemistry 2003; 278:20500–20506

52. Meyer EH, Giegé P, Gelhaye E, Rayapuram N, Ahuja U, Thöny-Meyer L, et al. AtCCMH, an essential component of the c-type cytochrome maturation pathway in *Arabidopsis* mitochondria, interacts with apocytochrome c. Proceedings of the National Academy of Sciences of the United States of America 2005; 102:16113–16118

53. Mand TD, Metcalf WW. Energy Conservation and Hydrogenase Function in Methanogenic Archaea, in Particular the Genus *Methanosarcina*. Microbiology and Molecular Biology Reviews 2019; 83:e00020–19

54. Wegener G, Krukenberg V, Riedel D, Tegetmeyer HE, Boetius A. Intercellular wiring enables electron transfer between methanotrophic archaea and bacteria. Nature 2015; 526:587–590

55. McGlynn SE, Chadwick GL, Kempes CP, Orphan VJ. Single cell activity reveals direct electron transfer in methanotrophic consortia. Nature 2015; 526:531–535

56. Scheller S, Yu H, Chadwick GL, McGlynn SE, Orphan VJ. Artificial electron acceptors decouple archaeal methane oxidation from sulfate reduction. Science 2016; 351:703–707

57. He X, Chadwick G, Kempes C, Shi Y, McGlynn S, Orphan V, et al. Microbial interactions in the anaerobic oxidation of methane: model simulations constrained by process rates and activity patterns. Environmental Microbiology 2019; 21:631–647

58. Doench JG, Fusi N, Sullender M, Hegde M, Vaimberg EW, Donovan KF, et al. Optimized sgRNA design to maximize activity and minimize off-target effects of CRISPR-Cas9. Nature Biotechnology 2016; 34:184–191

59. Kim SY, Ju KS, Metcalf WW, Evans BS, Kuzuyama T, van der Donk WA. Different biosynthetic pathways to fosfomycin in Pseudomonas syringae and Streptomyces species. Antimicrobial Agents and Chemotherapy 2012; 56:4175–4183

60. Metcalf WW, Zhang JK, Wolfe RS. An anaerobic, intrachamber incubator for growth of *Methanosarcina spp*. on methanol-containing solid media. Applied and Environmental Microbiology 1998; 64:768–770

61. Metcalf WW, Zhang JK, Apolinario E, Sowers KR, Wolfe RS. A genetic system for Archaea of the genus *Methanosarcina*: Liposome-mediated transformation and construction of shuttle vectors. Proceedings of the National Academy of Sciences of the United States of America 1997; 94:2626–2631

62. Boccazzi P, Zhang JK, Metcalf WW. Generation of dominant selectable markers for resistance to pseudomonic acid by cloning and mutagenesis of the ileS gene from the archaeon *Methanosarcina barkeri* Fusaro. Journal of Bacteriology 2000; 182:2611–2618

63. Deatherage DE, Barrick JE. Identification of mutations in laboratory-evolved microbes from next-generation sequencing data using breseq. Methods in Molecular Biology 2014; 1151:165–188

64. Gupta D, Sutherland MC, Rengasamy K, Mark Meacham J, Kranz RG, Bose A. Photoferrotrophs produce a PioAB electron conduit for extracellular electron uptake. mBio 2019; 10:e02668–19

65. Miller MA, Pfeiffer W, Schwartz T. Creating the CIPRES Science Gateway for inference of large phylogenetic trees. 2010 Gateway Computing Environments Workshop (GCE) 2010; 1–8

66. Chen IMA, Chu K, Palaniappan K, Pillay M, Ratner A, Huang J, et al. IMG/M v.5.0: An integrated data management and comparative analysis system for microbial genomes and microbiomes. Nucleic Acids Research 2019; 47:D666–D677

## Supplementary References

1. Kranz R.G., C. Richard-Fogal, J-S. Taylor, and E. R. Frawley, 2009 Cytochrome *c* biogenesis: Mechanisms for covalent modifications and trafficking of heme and for heme-iron redox control. Microbiology and Molecular Biology Reviews 73: 510–528. https://doi.org/10.1128/MMBR.00001-0

2. Mendler, K., Chen, H., Parks, D. H., Lobb, B., Hug, L. A., & Doxey, A. C. (2019). Annotree: Visualization and exploration of a functionally annotated microbial tree of life. Nucleic Acids Research, 47(9). https://doi.org/10.1093/nar/gkz246

3. Guss A. M., M. Rother, J. K. Zhang, G. Kulkarni, and W. W. Metcalf, 2008 New methods for tightly regulated gene expression and highly efficient chromosomal integration of cloned genes for *Methanosarcina* species. Archaea 2:193–203. https://doi.org/10.1155/2008/534081

4. Nayak D. D., and W. W. Metcalf, 2017 Cas9-mediated genome editing in the methanogenic archaeon *Methanosarcina acetivorans*. Proc. Natl. Acad. Sci. U. S. A. 114: 2976–2981. https://doi.org/10.1073/pnas.1618596114

